# Elucidating early intestinal stem cell response to bacterial infection

**DOI:** 10.1101/2024.09.15.613128

**Authors:** S. Lebon, N. Davidzohn, A. Habshush Menachem, A. Katz, N. Wigoda, R. Rotkopf, T. Dadosh, S. Levin-Zaidman, N. Dezorella, N. Blumberger, D. Hoffman, M. Hofree, M. Biton

**Affiliations:** Department of Immunology and Regenerative Biology, Weizmann Institute of Science, Rehovot, 76100, Israel; Bioinformatics Unit, Life Sciences Core Facilities, Weizmann Institute of Science, Rehovot 76100, Israel; Department of Chemical Research Support, Weizmann Institute of Science, Rehovot, 76100, Israel; The Lautenberg Center for Immunology and Cancer Research, Hebrew University of Jerusalem, Jerusalem 91120, Israel; The School of Computer Science and Engineering, Hebrew University of Jerusalem, Jerusalem 91904, Israel

## Abstract

Intestinal stem cells (ISCs) are the regenerative force of the gut epithelium. Lgr5+-ISC have been shown to respond to changes in their microenvironment by coping with different metabolites, adapting to caloric changes, and recovering from injury and inflammation. However, how pathogenic bacteria affect adult stem cell regeneration and, as a consequence, the overall tissue adaption to infection has yet to be explored in depth. Here, we interrogated early Lgr5+ ISC responses to an enteric intracellular pathogen by profiling individual IECs from the mouse small intestine. Utilizing GFP-labeled *Salmonella enterica,* we isolated intracellular invaded cells to elucidate invasion programs of epithelial cell subsets. In particular, we identified a *Salmonella*-specific infection signature comprised of antimicrobial peptide (AMP) genes, including the Defensin gene family. Our findings demonstrate that *Salmonella enterica* targets differentiated Paneth, enterocytes, and stem/progenitor cells at these early stages of infection. In response, a rapid Lg5+ ISC-driven cellular remodeling to enterocyte and Paneth lineages expressing AMP genes is initiated to combat the intruders. Importantly, we uncovered an ISC differentiation program via inflammasome activation to protect the crypt environment, while eliminating infected stem cells from the overall stem cell pool. This novel Lgr5+ stem cell defense mechanism not only protects the gut epithelium from persistent bacterial infection but also promotes tissue regeneration. We propose epithelial remodeling to AMP-secreting cells as a novel innate immune response to handle different gut stresses mediated by Lgr5+ ISCs to maintain organizational principles of gut homeostasis and physiology.

## Introduction

The small intestine functions to efficiently absorb nutrients and metabolites while simultaneously monitoring and responding to a variety of toxic substances or pathogens^1–3^. These responses are mediated by several distinct effector mechanisms: gut-resident immune cells, the enteric nervous system, local epithelial-specific defenses, and the secretion of gut hormones^4–8^. Like all mucosal tissues, the interface of the gut that directly contacts the lumen is a barrier comprised of epithelial cells, which act as the first line of defense^4,8–11^.

The cellular composition of the small intestinal (SI) epithelium is divided into two major lineages, absorptive and secretory^12^, reflecting its dual role as both a digestive tissue and a complex sensory-effector system. The gut epithelium of the SI is organized by a repeating villi structure, which projects toward the lumen, and invaginating crypts in which the stem and transit amplifying (TA) cells reside^13,14^. The crypt also contains Paneth cells, which secrete anti-microbial peptides or proteins (AMPs), such as defensins and lysozyme, into the lumen to further protect the crypt zone from bacterial penetration^15–17^. The mature IECs are rapidly differentiated from the stem cells to populate the epithelial barrier, and the overall turnover rate of the SI mature IECs is about 5 days, with the exception of Paneth cells ^13,16,18^. In contrast, intestinal stem cells (ISCs) are the only cell type that persists within the tissue. Consequently, it is imperative to protect ISCs from various gut stresses to prevent the initiation of gut pathologies. The stem cell niche within the crypt offers support and protection from the harsh lumen environment. The Lgr5+ ISCs at the base of the crypt are supported by epithelial, stromal, and immune cells, which provide essential growth factors essential for stem cell maintenance, such as the Wnt, Notch and EGF ligands^19–24^. In addition, Lgr5+ ISCs are also regulated by environmental factors, such as the microbiome and diet metabolites^25–29^, tissue injury^30–32^, and inflammation^33–37^ cues. However, how Lgr5+ ISCs respond to pathogenic bacteria remains poorly understood, representing a critical gap in our knowledge of stem cell biology.

Several previous studies have examined the gene expression profiles of individual IECs during pathogenic bacterial infection models, such as *Salmonella*^4^ or *Citrobacter*^11,38,39^, in the small and large intestines, respectively. The human version of these pathogens is prevalent worldwide and accounts for high rates of human morbidity and mortality^40^. *Salmonella enterica* serovar *Typhimurium* is a well-established bacterial model for enterocolitis and typhoid fever in mouse models^41,42^, and many studies investigated the pathophysiological changes in the course of these diseases^43–45^. However, an in-depth analysis of the Lgr5+ ISCs’ early responses to pathogenic bacteria is still required. In addition, our ability to distinguish between innate immune responses of Lgr5+ ISC compared to bystander stem cells under bacterial attack is still lacking.

Here, we used scRNA-seq to chart the early epithelial, particularly Lgr5+ ISC-specific responses to an intracellular bacterium, *Salmonella enterica*, in the small intestine. Using IEC single cell genomics and a GFP labeled *Salmonella* reporter, we generated an epithelial infection signature. Importantly, we found that many anti-microbial proteins and peptides, such as *Defa29* and *Lyz1*, and reduction of metabolic pathways, such as nucleotide metabolism, correlated with the *Salmonella enterica* invasion of epithelial cells. Finally, we characterized an intrinsic stem cell defense mechanism enabling infected stem cells to differentiate into AMP-secreting cells, mainly enterocytes, and Paneth cells, as a key stem cell response to the intracellular bacterium. This Lgr5+ ISC-specific inflammasome-mediated defense response allows rapid epithelial remodeling and helps to replenish the gut barrier while eliminating infected Lgr5+ ISCs and restoring gut homeostasis.

## Results

### Epithelial cell remodeling, an early response to *Salmonella* infection

The introduction of bacterial pathogens to the gut induces a strong inflammatory response to eliminate the pathogen from the intestine^46^. In the gut, the first responders to bacterial pathogens are the epithelial cells facing the lumen^47–49^. To investigate the Lgr5+ ISC and epithelial responses during early infection, we closely mimicked the natural route of *Salmonella enterica* infection without antibiotic pre-treatment (**Fig.1a**). We orally infected C57bl/6J mice with 5 × 10^8^ CFUs of *S. Typhimurium* (*S. Tm.*). SI IECs were collected at 0-, 12-, and 24 hours post-infection (*p.i*.). Notably, at 24h *p.i.*, we observed a marked increase in epithelial proliferation characterized by enlarged crypt length and elevation in Ki67+ cells, shown by combined single molecule fluorescence in situ hybridization (smFISH) and immunofluorescent staining (IFA) (**Fig. 1b** and **Extended Data Fig. 1a**). Despite the infection, no elevation in cell death was evident in these early time points, as assessed by terminal deoxynucleotidyl transferase dUTP nick-end labeling (TUNEL) assay, as well as the cleaved-Caspase 3 (CC3) staining (**Fig. 1b**, and **Extended Data Fig. 1b,e**) suggesting that the predominant cellular response at this stage is geared towards cell proliferation rather than cellular death or apoptosis.

**Figure 1:**
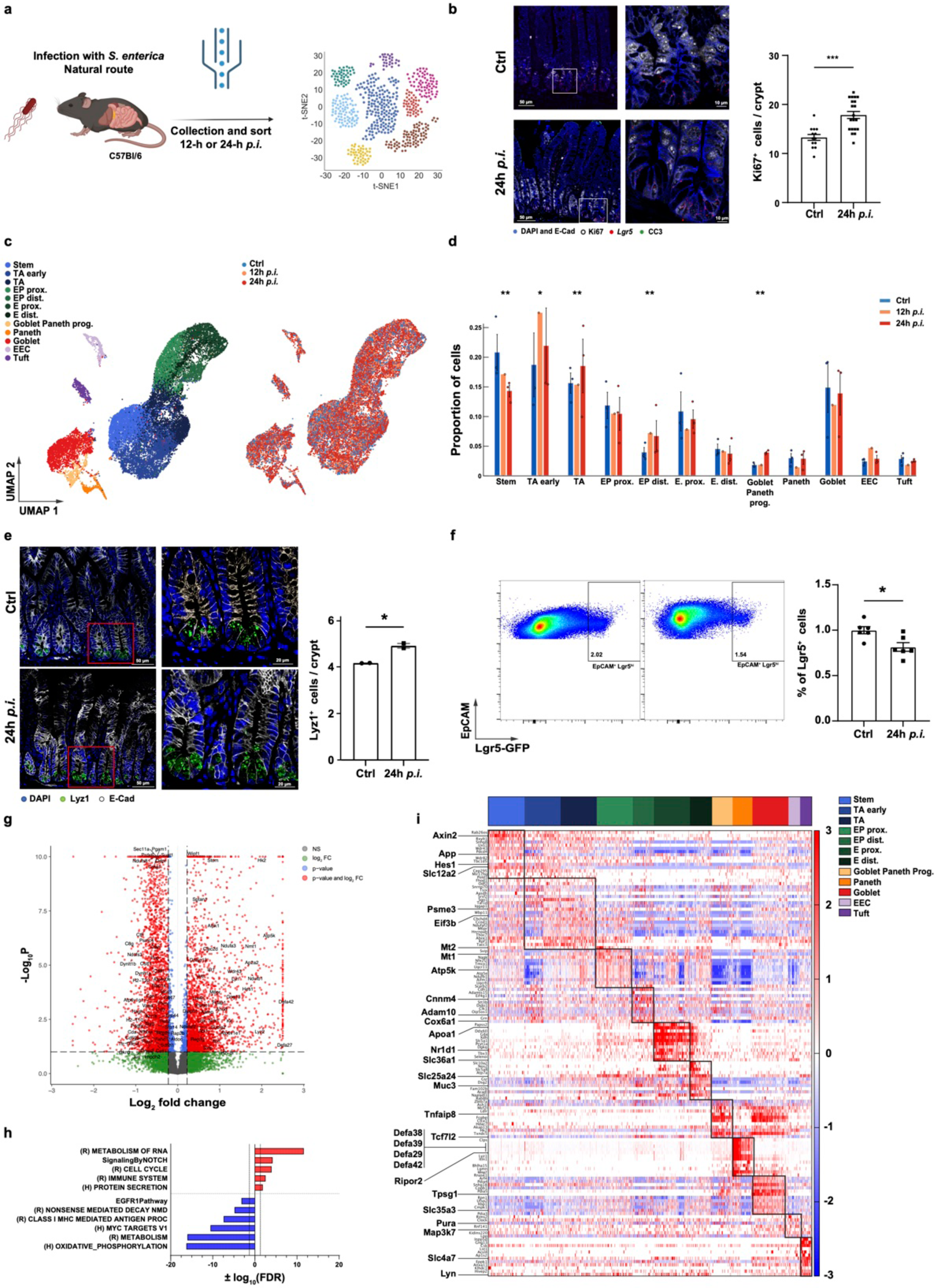
Early intestinal epithelial cell responses to *Salmonella enterica* infection. **a,** Experimental design. After per os infection by GFP-labelled *S. enterica,* intestinal epithelial cells (IECs) were extracted for downstream analysis. **b**, Representative images of immunofluorescence assay (IFA) and single molecule FISH (smFISH) of Ki67 (White), cleaved caspase 3 (CC3, green), E-cadherin (E-cad, blue), Lgr5 probe (red), and DAPI of SI sections from Ctrl and *Salmonella* 24h post-infection (*p.i.*). Scale bar 50 μm. Mid panel, insets 4x, scale bar, 10 μm. Right panel, number of Ki67+ cells per crypt was quantified; n ≥ 10 crypts per condition. Data are presented as mean ± SEM. ***p-value < 0.001, two-tailed Student’s *t-test*. **c**, Uniform manifold approximation and projection (UMAP) showing the 2D projection of single-cell RNA-seq epithelial cell profiles (*n* = 22,674 cells) colored by cell type (left) or condition (right). E, enterocyte; EEC, enteroendocrine; EP, enterocyte progenitor; TA, transit amplifying. **d**, Bar plots showing proportions of cell types for each mouse (points) and condition (FWER-Bonferroni adjusted \**p* < 0.05, \*\**p*< 8x10^-4^; Poisson glm comparing Ctrl vs 24h p.i.). Error bars, SEM. **e**, Representative image from IFA of Lyz1 (green) and E-Cadherin (white). Nuclei are shown using DAPI (blue) (left panel). The number of Lyz1+ cells per crypt was quantified (right panel); *n* = four mice analyzed; two-tailed Student’s *t-test*; data in graph are shown as mean ± SEM. \**p*-value < 0.05. **f**, Reduction in Lgr5+ ISC number 24 hours after *Salmonella* infection. Representative flow-cytometry (FC) plot (left) and quantification (right) of ISCs gated on EpCAM+ cells. *n* = 6 mice per condition; data in graph are shown as mean ± SEM; two-tailed Student’s *t-test* \**p*-value < 0.05. **g**, Volcano plot showing differentially expressed genes (DEG) in IECs 24h post-infection compared to control mice. Colored points correspond to false discovery rate (FDR) < 0.05 and/or log2(fold change) > 0.25. Color coding is shown on the top right. **h**, Selected gene sets showing hyper-geometric enrichment when testing the 400 significantly upregulated and downregulated genes in IEC 24h *p.i.* vs. control. H, Hallmark; R, Reactome. Plotted is the negative log10 of enrichment p-values. The direction of enrichment is shown as positive for gene-sets enriched in Sal24h p.i. and negative in ctrl. **i**, Cell-type specific gene expression in response to *Salmonella* infection 24h *p.i*. The color scale shows the expression z-score in 24h *p.i.* The cell subset bar is on top, and cell subset identifiers are on the top right.

To investigate the temporal alterations in the SI IECs in response to *S. Tm.* infection, IECs from infected or mock-treated mice were subjected to EpCAM+ sorting using flow cytometry (**Extended Data Fig. 1c**) followed by EpCAM+ hashed droplet-based 3’ scRNA-seq at 0-, 12-, and 24-hours *p.i.* using the 10X Chromium platform (**Methods**). We collected 26,094 cell profiles and retained 22,674 after excluding low-quality cells showing evidence of contamination (e.g., immune, doublets, etc.). Following filtering, we observed a median of 3,102 genes and 10,803 unique transcripts per cell. After normalization, variable gene selection, and dimensionality reduction, 12 clusters were identified by unsupervised graph clustering. Clusters were labeled by examination of differentially expressed genes (DEGs) and comparison to known marker genes **(Fig. 1c, Extended Data Fig. 1d,i** and **Supplementary Table 1**), consistent with recent reports^4–8,11,33,50,51^. Remarkably, we observed a dynamic, time-dependent remodeling of the cellular composition of the SI IEC following infection, with a significant decrease in stem cell proportions and an increase in progenitor cells, including transit amplifying (TA), enterocyte and Paneth progenitors (**Fig. 1d**). The increase in Paneth cells was validated by IFA for Lyz1, as well as in combined Periodic Acid Schiff (PAS)-Alcian Blue staining (**Fig. 1e** and **Extended Data Fig. 1g**). Moreover, we validated the reduction in Lgr5+ ISC numbers 24 hours *p.i.* by cytometry analysis of the Lgr5-GFP knock-in (KI) mouse model^52^ (**Fig. 1f**) and the stem cell marker, Olfm4 IFA (**Extended Data Fig. 1f**). In conclusion, we identified a rapid IEC remodeling mediated by stem cell reduction and higher proliferation at early time points of *Salmonella* infection.

Next, we examined the global transcriptional changes that occur 24 hours post-infection (*p.i.*) as well as the normalized data corrected for cell-type composition shifts resulting from cellular remodeling. Both approaches identified the upregulation of cell cycle and pro-inflammatory programs, including the epithelial-driven antimicrobial peptides and proteins, such as *Reg3g, Defa27, Defa42* and *Lyz2* genes^53–55^, while a reduction in metabolic processes, such as oxidative phosphorylation and MHC-I antigen processing pathway was evident (**Fig. 1g,h** and **Extended Data Fig. 1e,h and Supplementary Table 2**). These results suggested that epithelial cells already undergo cellular remodeling and a shift toward epithelial-specific innate immune transcriptional programs at this early time point.

We then defined *Salmonella*-induced gene expression for each cell subset using the scRNA-seq datasets, highlighting genes enriched in each particular subset (**Fig. 1i** and **Supplementary Tables 3–4**). While Paneth cells had elevated anti-microbial genes, such as *Lyz1* and *Defa29,* the Lgr5+ ISC had elevated expression of *Hes1*, a Notch signaling transcription factor associated with enterocyte differentiation^56^ and *Axin2*, a target of the Wnt pathway important for Paneth cell differentiation^20,57^ (**Fig. 1i**). These results indicate that Lgr5+ ISCs shift their transcriptional program to initiate a dynamic cellular remodeling toward Paneth and enterocytes.

Altogether, the early epithelial response to *Salmonella enterica* involves cell proliferation to minimize bacterial tissue barrier penetration, upregulation of AMP expression in all cells, and a rapid shift of differentiation toward professional AMP-secreting enterocyte and Paneth cells.

### Defined stem cell response to *Salmonella* invasion at single cell resolution

To interrogate the early response of Lgr5+ ISC invaded by *S.Tm.*, we exploited the intercellular invasion properties of *Salmonella*. We utilized GFP-labeled *Salmonella* to track invaded SI IEC (EpCAM+ GFP+) by flow cytometry analysis (**Fig. 2a**). We observed an increase in invaded IECs over time, rising from about 2.5% at 12 hours to nearly 7% at 24 hours *p.i*. (**Fig. 2b, Extended Data Fig. 2a**). To characterize the cell subsets that are invaded, we performed scRNA-seq of invaded EpCAM+GFP+ cells. We profiled 5,788 EpCAM+ GFP+ IECs 24 hours *p.i.* by comparing cell abundance within EpCAM+ and EpCAM+ GFP+ IECs, and identified *Salmonella+* (GFP+) enriched subsets (**Fig. 2c,d**; *n* ≥ 2 mice per group). Our analysis revealed that proximal enterocytes and Paneth cells were highly invaded by the pathogen as early as 24h *p.i*. (**Fig. 2d** and **Extended Data Fig. 2b,c**). Conversely, other mature secretory cell types, such as tuft, goblet, and enteroendocrine cells, had a low percentage of invaded cells (0.3%, 3%, and 0.4%, respectively).

**Figure 2:**
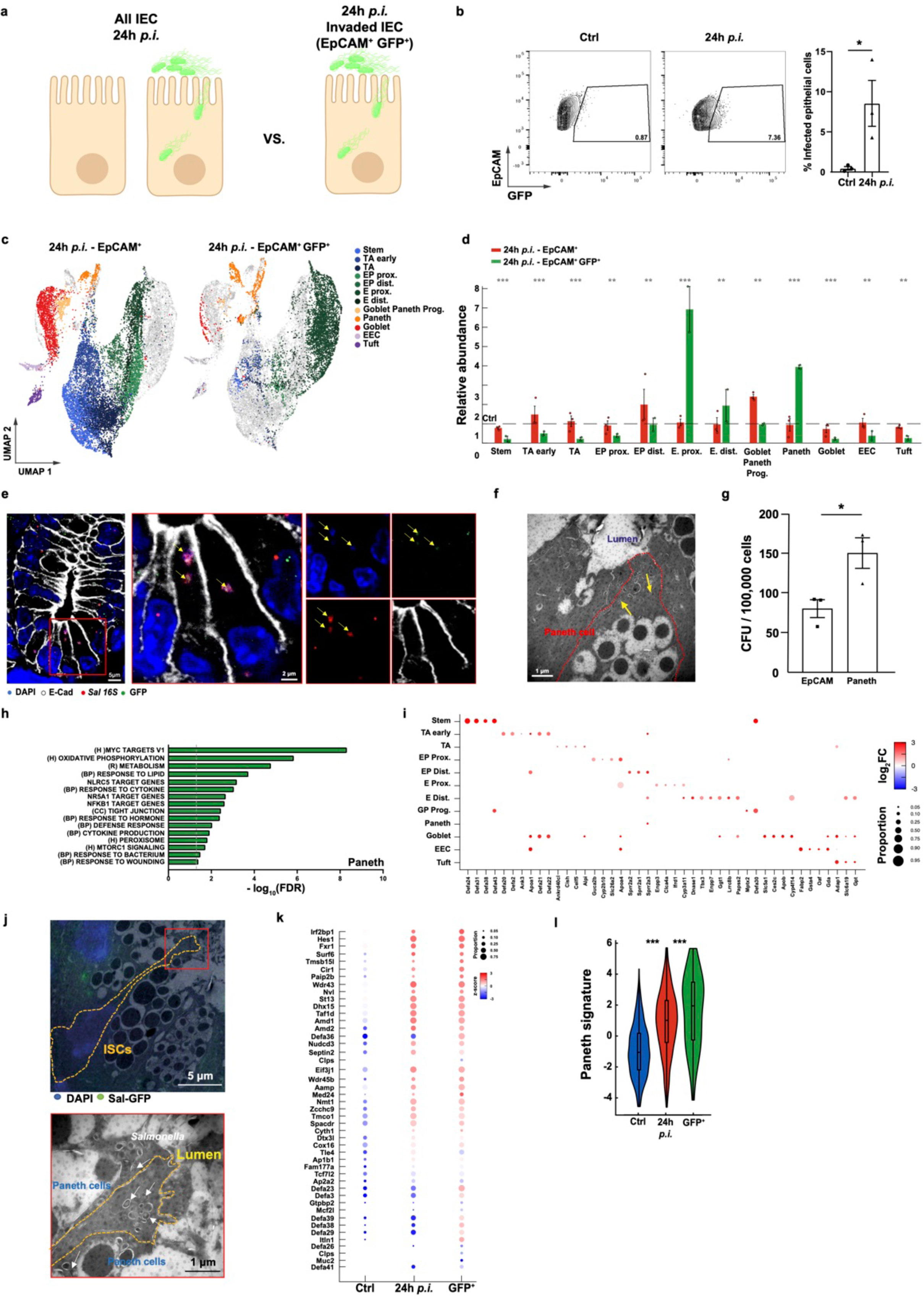
IEC-specific invasion responses. **a**, Experimental scheme. scRNA-seq analysis of invaded IEC (EpCAM+ GFP+) vs all IEC after 24h of *Salmonella* infection. **b**, Representative FC plot (left) and quantification (right) of invaded IECs (GFP+) or control (Ctrl) 24h p.i., *n* = 3 mice per condition. Data are presented as mean ± SEM. *p-value < 0.05, two-tailed Student’s *t-test*. **c**, UMAP presentation of scRNA-seq of epithelial cells colored by cell type (*n* = 18,266 cells) and divided by conditions. Left: 24h p.i. EpCAM+, right: 24h p.i. EpCAM+ GFP+. Grey dots represent the compared condition. EEC, enteroendocrine; EP, enterocyte progenitor; TA, transit amplifying. **d,** Cell-type relative abundance of IEC in 24 *p.i.* (red) and GFP+ cells (green) compared with control mean. Mean ratio ± SEM is depicted. *n*=3 mice. Statistical significance determined by Poisson-glm, stars indicate FWER-Bonferroni adjusted **P < 2.5x10-5. **e**, *Salmonella* invasion to cells at the bottom of the crypt. Left, smFISH of *Salmonella* 16S (red) co-stained with GFP (green) and DAPI (blue). E-Cadherin (grey) stains IEC boundaries. Scale bar, 5 μm. Mid-right, Inset, 3x; Scale bars, 2 μm. Right, single channel of each color. Yellow arrows, Triple-positive *Salmonella* particles. **f**, Transmission electron microscopy (EM) of *S. enterica* infected Paneth cell (white arrows). Dashed red line, Paneth cell boundaries; Scale bar, 1 µm. **g**, *Salmonella* invasion is enriched in Paneth cells. Colony-forming units (CFU) per 100,000 EpCAM+ or Paneth cells (EpCAM^+^ CD24^+^ FSC-A^high^). Data are presented as mean ± SEM of two independent experiments. \**p*-value < 0.05, two-tailed Student’s *t-test*. **h**, Selected gene sets showing hyper-geometric enrichment when testing the 400 significantly upregulated genes in invaded (GFP+) Paneth cells 24h *p.i.* vs. control. H, Hallmark; R, Reactome; BP, GO biological process; CC, GO cellular compartment. **i**, Dot plot of significantly differentially expressed genes (DEG) across cell-subsets comparing invaded (EpCAM+ GFP+) vs. EpCAM+ cells from control mice. Size of the dot indicates the frequency of cells, while strength of color indicates log2-fold change of mean expression (GFP+ vs Ctrl). Only dots indicating significantly expressed genes are shown. **j**, *Salmonella* particles in Lgr5+-ISCs. Upper panel, correlative light and electron microscopy (CLEM) of Lgr5+-ISC infected with *S. enterica* (GFP, green). Nuclei are shown using DAPI (blue). Yellow dashed line marks ISC boundaries; Scale Bar, 5 µm. Lower panel, magnification of the *Salmonella* particles. Inset: 4x; Scale bar, 1 µm; White arrows indicate *Salmonella*-GFP particles. **k**, Lgr5+-ISC-infection signature. Lgr5+-ISC-specific gene expression associated with response to *Salmonella* invasion (GFP+) or infection (24 *p.i*) compared to control stem cells. The size of the dot indicates the frequency of cells, while color indicates expression by Z-score of mean and std. **l**, Stem cell enrichment of Paneth cell signature under *Salmonella* infection. Paneth cell gene signature in control (ctrl), 24h *p.i.*, and GFP+ Lgr5+-ISC. Significance was determined by likelihood-ratio test of a generalized linear mixed-effects model (glme), \*\*\**p* < 5x10-9.

In *Salmonella*-infected cells, we noticed a four-fold enrichment in Paneth cells (**Fig. 2d** and **Extended Data Fig. 2b**), suggesting that *Salmonella* already reached the bottom of the crypt at 24 hours *p.i.*. To validate this result, we examined the presence of *S.Tm*. in the cells of the crypt zone at 24h *p.i.* and measured *Salmonella* colony forming units (CFU) from the whole crypt or dissociated crypt cells treated with 100 µg/mL Gentamicin for 30 minutes before seeding, to avoid the growth of attached extracellular *Salmonella* (**Extended Data Fig. 2d**). Next, we showed the presence of *Salmonella* in the crypt and Paneth cells *in situ* using IFA+smFISH of *Salmonella* GFP, and 16S rRNA analysis and electron microscopy, respectively (**Fig. 2e,f** and **Extended Data Fig. 2e**). Finally, we sorted EpCAM+ or Paneth cells from 24h *p.i.* crypts and found enrichment of *Salmonella* CFU in Paneth cells (**Fig. 2g**). *Salmonella* infection of Paneth cells was previously shown at 3 days *p.i.*^58^ and the secretion of the anti-microbial containing granules was reported^53,58^. Recent studies propose that Paneth cell hyper-secretion of AMPs might be beneficial to *Salmonella,* as a high concentration of AMP evacuates the microbiome niche and eases *Salmonella* penetration^59^, offering a possible explanation for *Salmonella*’s high invasion rate of Paneth cells.

Next, we explored the underlying IEC-specific invasion response at 24 hours *p.i.* (**Supplementary Tables 5-6**). Enterocytes and Paneth lineages, known for their antimicrobial defense systems^60,61^ upregulated genes associated with bacterial defense, including the bactericidal genes *Sprr2a1* and *2*^62^, and *Defa30* and *Mptx2* (**Fig. 2h,i** and **Extended Data Fig. 2f,g**). A more profound response of these cells involved the downregulation of metabolic processes involved in RNA and protein metabolism and translation (**Extended Data Fig. 2f**). Moreover, *Salmonella*-invaded Lgr5+ ISC exhibit a pronounced response to the invasion, including elevation of Defensin genes such as *Defa24* and *Defa31* (**Fig. 2i** and **Extended Data Fig. 2f,g**), suggesting a stem cell-mediated epithelial response integrating invasion and inflammation signals to *Salmonella* infection.

As mentioned above, the EpCAM+ GFP+ analysis showed a signal reflecting *Salmonella* invasion to Lgr5+ ISCs, which was accompanied by the induction of AMP genes (**Fig. 2i**). Lgr5+ ISCs are the regenerative compartment of the SI IECs, and infection with intracellular pathogens could be detrimental to the tissue. Lgr5+ ISC infection may result in persistent pathogen infection while reducing the tissue’s fitness to replenish the infected or damaged epithelial cells. However, our results suggested a quick turnover of the epithelial cells (**Fig. 1b-d**) and rapid cellular remodeling toward AMP-secreting cells in response to the pathogen (**Fig. 1d,e)**. Therefore, we further investigated the protective Lgr5+ ISC response to *Salmonella* invasion. First, we validated our single cell results by examining *Salmonella*’s invasion of stem cells using correlative light electron microscopy (CLEM) (**Fig. 2j**). Cytometry analysis and single cell data indicated a reduction in stem cell numbers under the early *Salmonella* infection (**Fig. 1d**), suggesting that stem cells either undergo cell death or rapid proliferation and differentiation. To distinguish between these possibilities, we compared the single cell gene expression of the invaded-(GFP+) versus unfractionated infected (24h *p.i.*) or control stem cells (Ctrl). We observed no change in apoptosis and an elevation in proliferation signatures in the infected stem cells (**Extended Data Fig. 2h**), as also validated by Ki67 and TUNEL staining (**Fig. 1b and Extended Data Fig. 1b**). In addition, we noticed a reduction in the ‘stemness’ signature of the *Sal*-invaded stem cells (**Extended Data Fig. 2h**). Examining *Salmonella*-ISC gene expression revealed an elevation in Wnt and Notch signaling, important for Paneth and Enterocyte lineage differentiation^20^, respectively (**Extended Data Fig. 2i**). Furthermore, we observed elevation in the expression of Paneth cell markers, mainly AMP genes such as those in the defensin family, differentiating between invaded (GFP+) and infected (24h *p.i.*) or control stem cells 24h *p.i.* (**Fig. 2i,k, Extended Data Fig. 2g**, and **Supplementary Table 7-8**). The expression of Paneth cell gene markers in invaded Lgr5+ ISCs prompted us to plot the Paneth cell gene signature in Lgr5+ ISCs under various conditions, including non-infected control, infected, or invaded ISCs. We identified significant upregulation of the Paneth cell program in the infected (24h *p.i.*) and invaded ISCs (GFP+) (**Fig. 2l**). These results support a novel Lgr5+ ISC-specific response to infection in which Lgr5+-ISCs either acquire the Paneth cell program to resist the intruder or differentiate toward the Paneth lineage.

### Charting early Lgr5+ ISC responses to *Salmonella* infection

To explore the two potential ISC decisions, either gaining the Paneth cell program or differentiation following early *Salmonella* infection (24h), we utilized the ‘scPulse-Seq’ method, combining scRNA-seq with lineage tracing of stem cells^63^. We generated an inducible *Lgr5*-driven *TdTomato* mouse strain (*Lgr5-CreER^T2^* crossed to *Rosa26*-*LSL-tdTomato*) to label Lgr5+ ISCs and their progeny (**Fig. 3a**). *tdTomato* expression was induced immediately after infection with *S. enterica*, and the generation of newly differentiated cell types was monitored (**Fig. 3b**). Live EpCAM+ tdTomato+ cells were isolated 24 hrs after tamoxifen (TAM) injection from intestinal crypts of control or infected (24h *p.i.* with *S.tm*) cells and subjected to scRNA-seq (**Extended Data Fig. 3a**). After quality control, our dataset encompassed a total of 14,068 cells with an average detection of 1,792 genes per cell. We identified 14 distinct epithelial subsets along the differentiation axis (including stem cell subsets^33^ and early progenitors, and assigned unique cell-subset identities to each cluster (**Fig. 3c, Extended Data Fig. 3b**, **Supplementary Table 9**, *n* = 2 mice per condition). Cell subset proportion analysis between *Salmonella*-infected and control showed a massive reduction in ISC subsets, ISC-I and -II, and a mild reduction in ISC-III, while a significant expansion of the Paneth subsets was observed (**Fig. 3d**). TdTomato+-labeled cells across each major cell subset revealed a faster IEC turnover and reduced precursor cells with the exception of Paneth progenitors. Conversely, populations of differentiated cells expanded following infection, particularly noticeable in both enterocytes and Paneth cells, while goblet and tuft cells did not change in their proportions, and EECs were absent from the analysis (**Fig. 3d**). This phenomenon is highlighted along the differentiation continuum by a diffusion map^64^ of all tdTomato+ cells from the infected mice compared to the non-infected controls, emphasizing direct differentiation from stem cell to Paneth lineage, as was suggested before^65^ (**Fig. 3e)**. Next, we applied trajectory inference using Monocle 3^66^ and constructed the absorptive- and secretory-lineage trajectories. We visualized the predicted differentiation trajectories of Lgr5+ ISCs using pseudotime alignment (**Extended Data Fig. 3c,d**). The secretory pseudotime trajectory suggested a reduction in stem cell states and elevation in mature Paneth cell subsets 24h *p.i.* by accelerating the differentiation of Paneth cell progenitors directly from the stem cells (**Extended Data Fig. 3c**). The enterocyte lineage differentiation was elevated as well (**Extended Data Fig. 3d)** in concordance with our previous analysis, in which enterocyte and Paneth cell lineages were shown to be elevated upon *S.Tm* infection (**Fig. 1d**). Inspection of Gene Set Enrichment Analysis (GSEA) or gene ontology (GO) analysis of all ISC subsets identified mTOR pathway enrichment, which could account for Paneth cell differentiation, as described before^67^ (**Fig. 3g and Extended Data Fig. 3f**). Next, qPCR of newly differentiated (EpCAM+ tdTomato+) cells confirmed the decrease in Lgr5+ ISC marker expression, alongside increased expression of Paneth cell markers in the infected distal SI (**Extended Data Fig. 3e**). Furthermore, an increase of Paneth cells relative to the total population of newly formed IECs was observed by flow cytometry analysis (**Fig. 3h**). This trend was absent in mice infected with Heat-Killed *Salmonella*, suggesting the importance of cell invasion to this phenomenon (**Extended Data Fig. 3g**). These results support a rapid cellular remodeling of IECs, especially a turnover of ISC to Paneth cell lineage 24h *p.i*.

**Figure 3:**
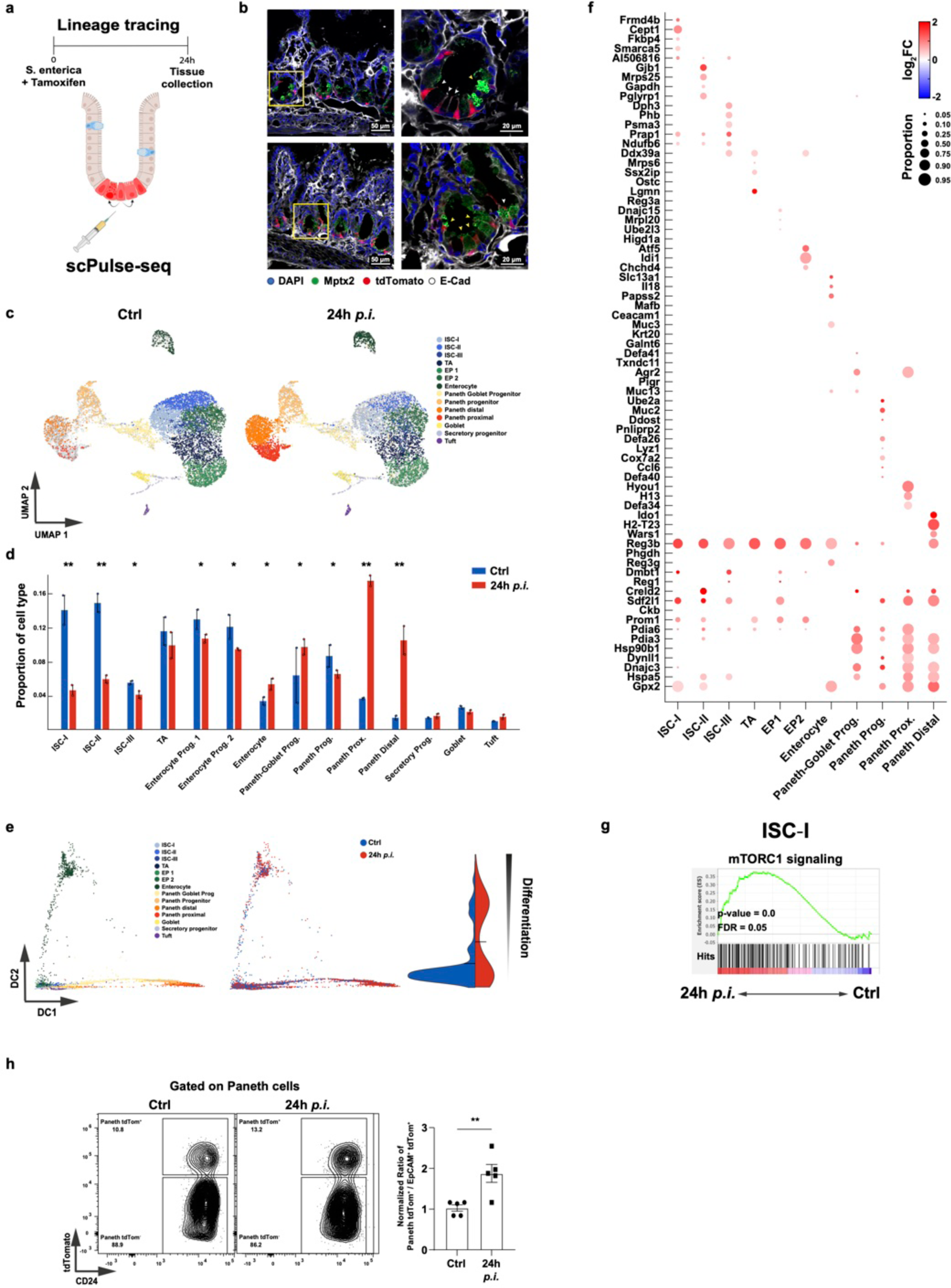
Lineage tracing of IEC differentiation during *Salmonella enterica* infection. **a,** Experimental scheme of IEC fate mapping during Salmonella using Lgr5-CreER^T2^:LSL-tdTomato mouse model. Mice were infected with GFP-labelled S. enterica in parallel to Tamoxifen injection to induce tdTomato expression in Lgr5+-ISCs and their progeny. **b**, Representative images of ISC-driven tdTomato from Ctrl and 24h *p.i..* Distal intestine was stained for E-Cadherin (white), Mptx2 (green), DAPI (blue) and tdTomato (red, endogenous); Scale bar, 50 µm. Right, insets showing higher magnification of boxed areas. White arrowheads, ISC tdTomato+; yellow arrowheads, Paneth cells tdTomato+; Scale bar 20 µm. **c**, UMAP of TdTomato+ IECs (14,068 cells) colored by cell subsets and divided by condition (left: control, right: 24h *p.i.*). Grey dots, depicting the cells from the other conditions; *n* = 2 mice per condition. ISC, intestinal stem cells; TA, transit amplifying; EP, enterocyte progenitor. **d,** Differentiation shift under *Salmonella* infection. Cell type proportions under the different conditions, control (blue) and infected (red). Cell type proportions for each mouse (points, *n*=2 mice per condition) and condition (FWER-Bonferroni adjusted \**p* < 0.05, \*\**p*<0.004; Poisson glm comparing Ctrl vs +24h *p.i.*). Error bars, SEM. **e**, Diffusion maps (Destiny; **Methods**) of tdTomato+ epithelial cells colored by cell subset (left) or condition (center). Right, a diffusion pseudotime (dpt) score comparing the differentiation state of control and 24h *p.i.* **f,** Dot plot of significantly differentially expressed genes (DEG) across cell subsets comparing EpCAM+ tdTomato+ cells from control or 24 *p.i.*. Size of the dot indicates the frequency of cells, while color indicates log2-fold change of mean expression between the conditions. Only dots indicating significantly expressed genes are shown. **g**, Enrichment of the mTORC1 signaling pathway in ISC-I at 24h *p.i*. using gene set enrichment analysis (GSEA). FDR, false discovery rate. **h,** Representative flow cytometry plot of the newly differentiated Paneth cells (Paneth tdTom+) (left), and quantification of the normalized ratio of Paneth tdTom+ compared to all tdTomato+ EpCAM+ (EpCAM+ tdTom+) cells (*n* = 5 mice per condition). \*\**p*-value < 0.01, two-tailed student’s *t-test*.

### Intrinsic stem cell defense mechanisms to bacterial infection

To assess if immune cells contribute to the cellular remodeling of Lgr5+ ISC at early time points (24h), we utilized the TKO immunodeficient mouse model with defective expression of the recombination activating gene 2 (Rag2), the CD47 antigen gene, and the X-linked interleukin 2 receptor, gamma chain (Il2rψ), lacking both functional innate and adaptive immune system. Overall, the number of Paneth cells in TKO mice was reduced compared to WT mice, but an increase in Paneth cell number 24h *p.i.* was still observed, similar to that seen in immunocompetent mice (**Extended Data Fig. 4a,b**). These results suggested that stem cell-guided differentiation toward Paneth cells at this early time point is an intrinsic cellular process. To examine the direct Lgr5+ ISC intrinsic responses to a pathogenic bacterium, we utilized an *ex vivo* organoid system^68,69^. We isolated distal crypt from WT mice and infected them with *Salmonella* at a multiplicity of infection (MOI) of 10 bacteria per cell for 2 hours. The organoids were cultivated with Gentamicin for an additional 24 hours. qPCR analysis showed a decrease in Lgr5+-ISC gene markers, as was reported before^70^, and an increase in Paneth cell gene markers compared to the uninfected control (**Fig. 4a-c**). In addition, using flow cytometry, we observed an increase in the proportion of Paneth cells identified by either CD24 expression and high forward scatter (FSC) or Lyz1+ staining^71^ (**Extended Data Fig. 4c,d**). Next, to verify a direct Lgr5+ ISC involvement, we sorted stem cells from the Lgr5-GFP mouse model^52^ and infected them with *S.Tm.* (MOI of 10) for 2h. After 24 hours, cells were collected, and qPCR and flow cytometry analysis were conducted (**Fig. 4d-f and Extended Data Fig. 4e**). Paneth cell differentiation was elevated, indicating the activation of an intrinsic Lgr5+ ISC-specific defense mechanism. Next, we assessed whether Paneth cell differentiation was associated with a more robust antibacterial response by testing the ability of *Salmonella* to survive within Paneth cells compared to Lgr5+ ISC. To this end, we infected sorted Paneth or Lgr5+ ISC with *S.Tm*. for 2 hours (MOI of 10) and examined *Salmonella* survival within each of these cell types. While both Lgr5+ ISC and Paneth cells were invaded by *Salmonella* after 2 hours of incubation, *Salmonella* survived after 12h within Lgr5+ ISC but not in Paneth cells (**Fig. 4g**). After 24h, neither Paneth nor Lgr5+ ISC had live bacteria within the cells (**Fig. 4g** and **Extended Data Fig. 4f**). These results suggested the presence of an Lgr5+ ISC-specific defense mechanism inducing differentiation towards Paneth cells, which are more effective at eliminating *Salmonella* invasion to avoid persistent infection.

**Figure 4:**
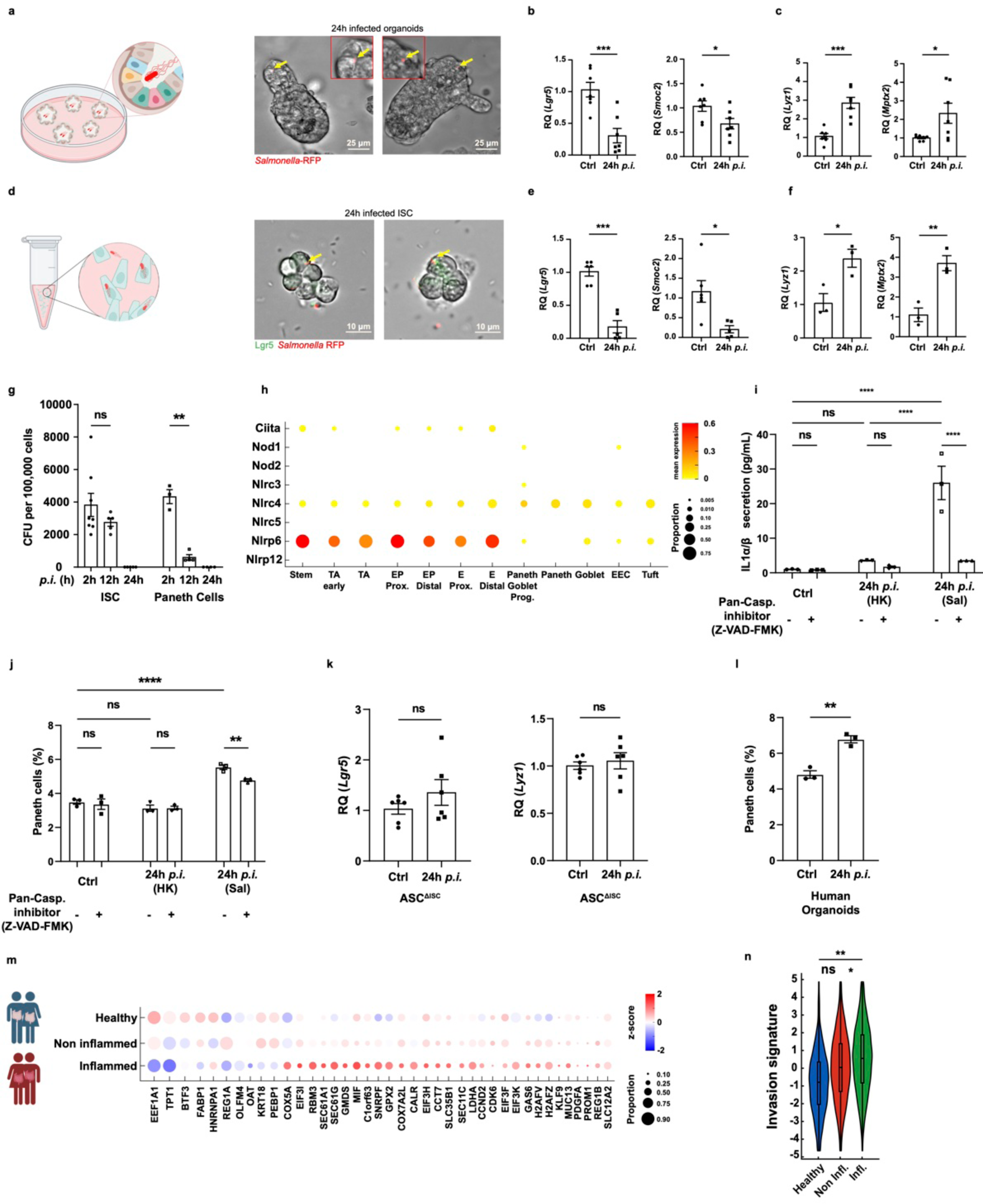
Stem cell-intrinsic defense mechanisms against *Salmonella* infection. **a**, Organoid infection scheme (left). Representative pictures of infected organoids were taken 24h *p.i.* (right). Invaded cells are marked by a yellow arrow (RFP-labeled *Salmonella)*; Scale bar, 25 µm. **b-c**, Relative quantification (RQ) of Lgr5+-ISC markers, *Lgr5* (**b**, left) and *Smoc2* (**b**, right) or Paneth cell markers *Lyz1* (**c**, left) and *Mptx2* (**c**, right) in control compared to *Salmonella*-infected organoids, 24-hours *p.i*. using quantitative PCR (qPCR). *n*=7 biological repetitions, \**p*-value <0.05, \*\*\**p*-value < 0.001, two-tailed student’s *t-test.* **d**, Stem cell infection scheme (left). Representative pictures of RFP-*Salmonella* invaded ISCs were taken 24h *p.i.* (right); Scale bar, 10 µm. **e-f**, Relative quantification (RQ) of Lgr5+-ISC markers, *Lgr5* (**e**, left) and *Smoc2* (**e**, right) or Paneth cell markers *Lyz1* (**f**, left) and *Mptx2* (**f**, right) in control compared to *Salmonella*-infected Lgr5+-ISC, 24-hours *p.i*. using quantitative PCR (qPCR). *n*≥3 biological repetitions, \**p*-value <0.05, \*\**p*-value < 0.01, \*\*\**p*-value < 0.001, two-tailed student’s *t-test.* **g**, CFU per 100,000 cells (ISC or Paneth cells) after 2-, 12- or 24-h of *in vitro* infection. Data are presented as mean ± SEM of two independent sets of experiments. *n*≥3 biological repetitions; ns, non-significant, \*\**p*-value < 0.01, one-way ANOVA. **h**, Dot plot of NOD-like receptors expressed in different IECs in homeostasis. Size of the dot indicates the proportion of cells expressing a gene, while color indicates mean expression level. **i**, Levels of interleukin-1α or β (pg/ml) present in the media of control, heat-killed (HK) *Salmonella* or *Salmonella*-RFP infected ISC 24h *p.i.,* with or without pan-caspase inhibitor (Z-VAD-FMK). Data are presented as mean ± SEM; ns, non-significant, *n*=3 biological repetitions, ****p-value < 0.0001, one-way ANOVA. **j**, Frequencies of CD24+ FSC-A^high^ cells (Paneth cells) derived from control ISC cells (ctrl) or challenged with HK Salmonella or Salmonella-RFP (*n* = 3 biological repetitions) tested by FC. Data are presented as mean ± SEM; ns, non-significant, **p-value < 0.01, ****p-value < 0.0001, one-way ANOVA. **k,** Relative quantification (RQ) of ISC marker, *Lgr5* (Left) and Paneth cell marker, *Lyz1* (Right) in control or *Salmonella* infected ASC^ΔISC^ organoids, 24-hours *p.i.*; ns, non-significant, two-tailed student’s *t-test.* **l,** Frequencies of Paneth cells derived from control human organoids cells (ctrl) or challenged with *Salmonella*-RFP (*n* = 3) tested by FC. Data are presented as mean ± SEM \*\**p* value < 0.01, two-tailed Student’s *t-test*. **m,** Dot plot of selected genes from consensus *Salmonella* invasion signature, derived from shared differentially expressed genes (DEGs) across all experiments. The size of the dot indicates the frequency of cells, while color indicates expression by Z-score of mean and std. **n,** Enrichment of consensus *Salmonella* invasion signature in inflamed ISCs of CD patients^50^. Significance is determined by likelihood ratio test; ns, not significant; \**p* value < 0.05, \*\**p* value < 0.01.

It is well established that intracellular bacteria can trigger the inflammasome, the innate immune sensor, operated via the Nod-like receptors (NLRs)^72–74^. Indeed, an inspection of NLR members in Lgr5+ ISC using the IEC single cell data revealed that many NLR members are expressed by Lgr5+ ISCs, including high expression of the epithelial enriched *Nlrp6*^72,75^(**Fig. 4h**). To examine the involvement of the inflammasome in such an ISC-specific defense mechanism, we infected sorted Lgr5+ ISC with either live or heat-killed *Salmonella* for 2h (MOI of 10) (**Extended Data Fig. 4g**). After 24 hours, we transferred the supernatant to HEK-Blue interleukin 1α/μ (IL1α/μ) or IL-18 reporter cells to examine inflammasome activation following *Salmonella* infection. Infected Lgr5+ ISCs secreted high levels of IL-1α/μ and IL-18 following infection compared to infected Paneth cells (**Extended Data Fig. 4h**). Next, to examine the inflammasome involvement in Paneth cell differentiation, we inhibited the inflammasome activation by the pan-caspase inhibitor (Z-VAD) and tested either IL-1α/μ secretion from the Lgr5+ISC using the HEK-blue reporter cells or Paneth cell percentage by flow cytometry. Reduced IL-1α/μ secretion was observed from Lgr5+ ISCs, and lower Paneth cell percentage was determined in the infected Lgr5+ ISCs treated with Z-VAD (**Fig. 4i,j**). Finally, we genetically inhibited the inflammasome pathway by depleting the adaptor protein of inflammasome activation, the *Pycard* (ASC) gene, in IECs by generating ASC^ΔISC^ mice. Organoids from ASC^ΔISC^ infected with *Salmonella* (MOI of 10) showed no elevation in the Paneth cell marker, *Lyz1*, nor a reduction in the stem cell marker, *Lgr5*, tested by qPCR (**Fig. 4i**). These results support the activation of the inflammasome in Lgr5+ ISCs, which promote differentiation processes upon intracellular bacterium invasion to avoid persistent Lgr5+ ISC infection and eliminate the bacteria as part of their differentiation to an AMP-secreting Paneth cell.

Our findings show that murine Lgr5+ ISC responds to *Salmonella* infection by differentiation to Paneth cell lineage. To examine if this Lgr5+ ISC response to Salmonella exists in humans, we utilized human ileal *ex vivo* organoid model. We infected the organoids with *Salmonella* at a MOI of 10 for 2h. 24h post-infection, we observed an increase in Paneth cell proportions, in accordance with the mouse model (**Fig. 4l**). Next, we isolated enriched stem/progenitors from dissociated early organoids (**Methods**) and infected them with *S.Tm.* (MOI of 10) for 2h. 24h later, we observed a high proportion of Paneth cells from the *S.Tm*. infected cultures, further supporting the activation of this Lgr5+ ISC-specific defense mechanism in humans (**Extended Data Fig. 4j**). Patients with inflammatory bowel disease (IBD) are at an increased risk for infections, including a heightened susceptibility to *Salmonella*^76,77^. We tested the *Salmonella* infection signature on published single cell data of terminal ileum from Crohn’s disease (CD) patients, focusing on ISC from inflamed, non-inflamed, or healthy individuals^50^. We constructed a consensus *Salmonella* invasion signature derived from the differentially expressed genes identified across our various experimental datasets (**Fig. 4m**). Our results support the enrichment of the *Salmonella*-infected ISC signature in inflamed ISC from CD patients, while a trend but no significant enrichment was found in non-inflamed tissue (**Fig. 4n**). These results link pathogenic bacterial infections or Lgr5+ ISC defense mechanism to intestinal flares in IBD patients, suggesting that further investigation of the Lgr5+ ISC innate immune response in recurrent IBD flares is warranted. Overall, the direct Lgr5+ ISC defense mechanism to bacterial infection described here (**Extended Data Fig. 4k)** may have further implications for gut health in homeostasis and disease.

## Discussion

Adult stem cells are the regenerative compartment of tissues coordinating tissue renewal and contraction. The intestinal stem cells, the Lgr5+ ISCs, are highly proliferative stem cells responsible for replenishing the intestinal epithelial cells (IEC) to maintain the monolayer epithelial barrier, generating cell types with absorptive or secretory properties to execute the needs of the tissue. Therefore, tight regulation of Lgr5+ ISC signals is provided by Paneth cells^19,20^, stromal^21–24^, and immune cells^33,78^. In addition, external signals derived from microbial and diet factors have a critical role in maintaining Lgr5+ ISC^25,27,29^. However, how stem cells directly respond to pathogenic bacteria in the gut is less explored.

*Salmonella* infection is a common gastrointestinal bacterial illness, causing significant human morbidity and mortality globally. It is estimated to cause over 150,000 deaths annually, with 85% of *Salmonella* cases being food-related^79^. Therefore, it is important to attain a better understanding of the pathophysiological processes associated with *Salmonella* infection in the context of Lgr5+ ISCs. To examine the role of Lgr5+ ISC in defending against *Salmonella*, we utilized single-cell RNA sequencing to map the initial responses of the epithelial cells to the intracellular bacterium *Salmonella enterica* in the small intestine. These experiments were performed without antibiotic treatment to closely mimic a natural, spontaneous infection. To this end, we leveraged the *Salmonella* intracellular environment by employing GFP labeling of the infectious bacterium to facilitate isolation and the identification of invaded IECs from the small intestine. A single-cell analysis of GFP+ IECs was instrumental in elucidating the early, epithelial-specific responses to *Salmonella enterica*, and revealed enrichment of Paneth and proximal enterocytes in response to infection. We identified a global epithelial elevation in innate defense genes, Lgr5+ ISC induction of the Wnt, Notch, and mTOR signaling pathways resulting in rapid cellular remodeling of the gut epithelium associated with early *Salmonella* infection. Importantly, we found upregulation of anti-microbial peptides, including *Reg3b* and *Reg3g*, expressed specifically by enterocytes under steady state^4,80^ and known to be crucial for preventing attachment of bacteria to the epithelium^81^, and Defensin genes constitutively expressed by Paneth cells^82^. These findings indicate that some level of epithelial cell-type identity is lost under this pathological condition to induce anti-microbial defense.

Lgr5+ ISCs compete for space at the bottom of the crypt via neutral drift and symmetrical cell divisions to maintain stem cell numbers in homeostasis^83,84^. Here, we characterized an intrinsic Lgr5+-ISC defense mechanism triggering bacterial-infected stem cells to differentiate into AMP-secreting cells, mainly enterocytes, and Paneth cells, as a key stem cell response to the intracellular bacterium. This differentiation process is driven by inflammasome activation to eliminate infected stem cells and strengthen gut barrier restoration while elevating the fitness of the epithelial cells to combat the intruder. We validated our results *in vivo* using single cell Pulse-seq (scPulse-seq)^63^ to trace enterocyte and Paneth cell differentiation from Lgr5+ ISCs, and by *ex vivo* organoid or stem cell infection models under inflammasome inhibition or depletion. scPulse-seq and the organoid or stem cell *ex vivo* experiments showed a substantial cellular remodeling of the gut SI epithelium, initiated by Lgr5+ ISC differentiation, mainly toward Paneth cells under early bacterial infection. The differentiation of stem cells in response to infection increases Lgr5+ ISC readiness to initiate IEC adaption to the hostile environment of the gut.

Our research indicates that the stem cell defense response to bacterial infections is not limited to mouse Lgr5+ ISCs. We show an elevation in Paneth cell differentiation in human terminal ileum organoids or stem/progenitor cells challenged with *S.Tm*., suggesting that the Lgr5+ ISC bacterial defense mechanism is conserved. In addition, examining the *Salmonella*-invaded ISC signature in Crohn’s disease (CD) patients single cell dataset^50^, we found a statistical increase of the signature in inflamed ISCs compared to non-inflamed or healthy ISCs. These findings suggest that the Lgr5+ ISC bacterial defense mechanism might be involved in intestinal flares in IBD patients, and further examination of this mechanism in the context of IBD is warranted.

In this study, we demonstrate how IEC cell types and states change dynamically as the small intestine adapts to infection by the intracellular pathogen. Our comprehensive, high-resolution map of early *Salmonella enterica* infection highlights how changes in epithelial cellular remodeling mediated by the Lgr5+ ISC innate immune response can play a key role in response to pathogens. The Lgr5+ ISC response to infection could have significant implications for understanding human gut health, stem cell biology, and potential therapies for colorectal cancer and IBD.

## Acknowledgments

We thank Prof. Ramnik Xavier (Broad Institute) and Prof. Roi Avraham (WIS) for their critical comments and discussions. We thank Shally Schwarzbaum for editing the paper, Merav Kedmi for help in RNA single-cell experiments, and the histology unit. M.B. holds the Ernst and Kaethe Ascher Career Development Chair. This study was supported by research grants from the Center for New Scientists at the Weizmann Institute of Science, the Israel Science Foundation (grant No. 1587/20), the Helen and Martin Kimmel Institute for Stem Cell Research at The Weizmann Institute of Science, and the Minerva Foundation, with funding from the Federal German Ministry for Education and Research, the Moross Integrated Cancer Center, the Israel Ministry of Science (IMOS, Grant No. 4631), the Dr. Gilbert S. Omenn and Martha A. Darling Weizmann Institute - Schneider Hospital Fund for Clinical Breakthroughs through Scientific Collaborations, a research grant from the Snider Foundation, the Abisch-Frenkel RNA Therapeutics Center, a research grant from the Shimon and Golde Picker, and a research grant from the Herbert K. Bennett Charitable Fund and Dwek Institute for Cancer Therapy Research.

## Author contributions

S.L. and M.B. conceived the study, designed experiments and interpreted the results; M.H. designed and performed the computational analysis with the assistance of N.B.; N.W. and R.R. performed the analysis of the lineage tracing experiment; M.B. and M.H. supervised this study; S.L. carried out all experiments; N.D. assisted with single cells experiments; S.L., S.L.Z., T.D. and N.D. performed the CLEM experiments; A.H.M. and A.K. assisted with immunohistochemistry and immunofluorescence experiments; D.H. assisted with infection experiments; and S.L., M.H., and M.B. wrote the manuscript, with input from all authors.

## Extended Data Figures

**Extended Data Figure 1:**
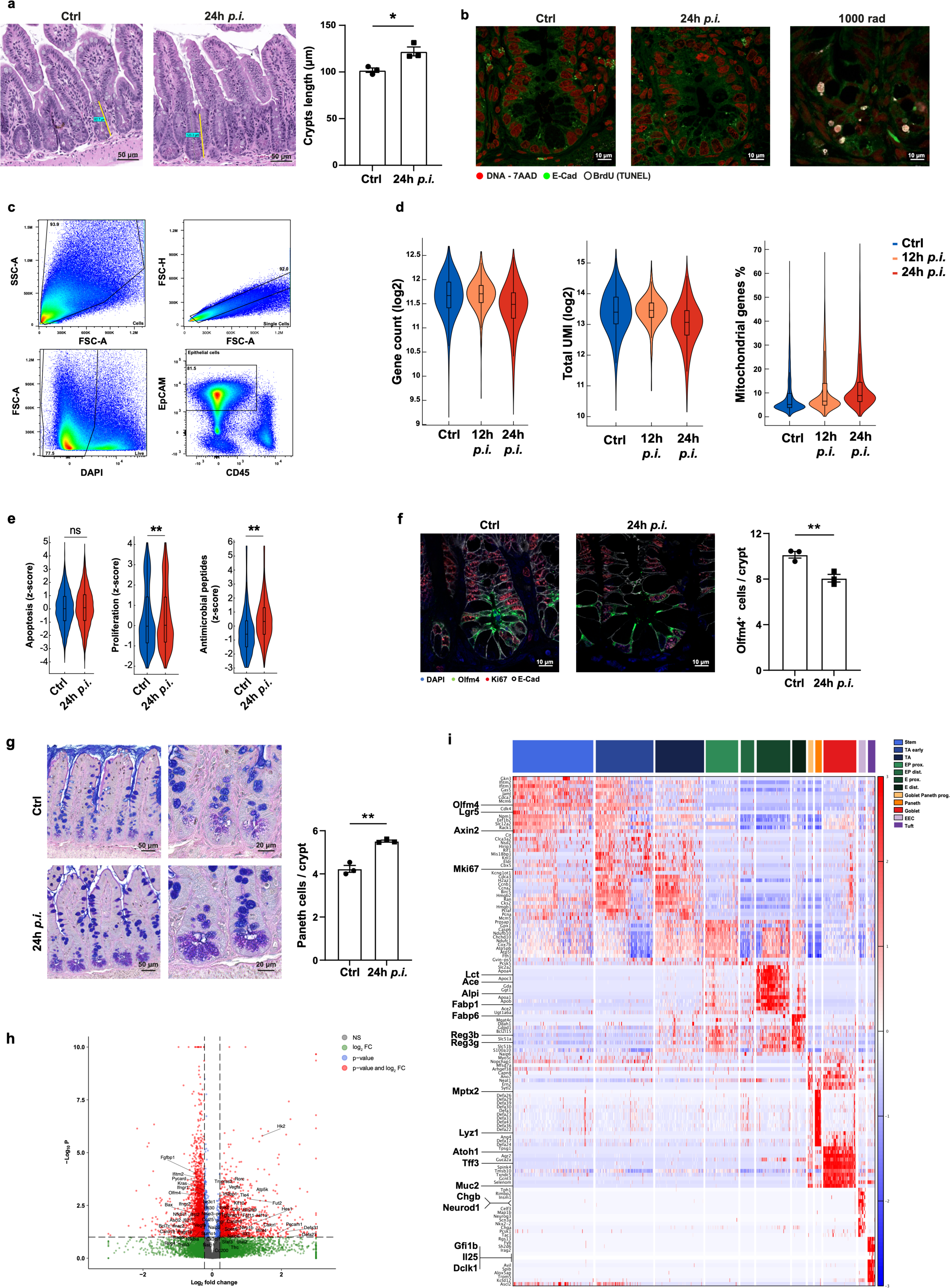
scRNA-seq quality control and subset annotations. **a,** Images of ileum sections from control and 24h *p.i.* mice using H&E staining, scale 50μm. Crypt length was quantified (right); *n* = 3 mice analyzed per group, 3 independent experiments; data are presented as mean ± SEM \**p* value < 0.05, two-tailed Student’s *t-test*. **b,** Representative images of ileal crypts staining of terminal deoxynucleotidyl transferase dUTP nick-end labeling (TUNEL) assay, BrdU (white), DNA – 7AAD (red) and E-Cadherin (E-cad, green) from Ctrl, *Salmonella* 24h post-infection (*p.i.*) and 24h after irradiation (1000 rad) mice. Scale 10μm. **c,** Flow cytometry gating strategy of IECs sorting for live EpCAM^+^ CD45^-^ cells. Indicated numbers are frequencies from parental population. **d,** Cell quality metrics for scRNA-seq data. Distributions of the number of genes detected with non-zero transcript counts per cell (left), the number of total unique molecular identifiers (UMI) detected per cell (center), and the fraction of reads mapping to mitochondrial genes per cell (right) are shown. **e,** Distribution of the expression of apoptosis, proliferation, and antimicrobial peptides (AMP) pathway genes in intestinal epithelial cells from control (ctrl) or 24h *p.i.*; ns, not significant, **FDR < 0.01; *Mann–Whitney U test*. **f,** Immunofluorescence assay (IFA, left) and quantification (right) of Olfm4 (green), Ki67 (red), and E-Cadherin (E-cad, white). Nuclei are shown using DAPI (blue). Scale 10μm. The number of Olfm4^+^ cells per crypt was quantified. *n* = 3 mice per group; data are mean ± SEM; \**p*-value < 0.05, two-tailed student’s *t-test*. **g,** Periodic Acid Schiff (PAS) and Alcian Blue (AB) staining of ileum sections from control and 24h *p.i.* mice, scales 50μm (left) and 20μm (mid). The number of Paneth cells per crypt was quantified (right); *n* = 3 mice per group; data are mean ± SEM, \*\**p*-value < 0.01, two-tailed student’s *t-test*. **h,** Volcano plot showing differentially expressed genes (DEG) in IECs 24h *p.i.* compared to control mice, adjusted to account for differences in cell-type distribution between the conditions by cell sub-sampling (**Methods**). Colored points correspond to false discovery rate (FDR) < 0.05 and/or log2(fold change) > 0.25. Legends plotted on the top right. **i,** Cell subset signatures. The normalized expression level (color scale, row-wise z-score of expression) of genes (rows) across cells (columns) is shown sorted by cell subset. The cell subsets bar is on top, and legends are on the top right.

**Extended Data Figure 2:**
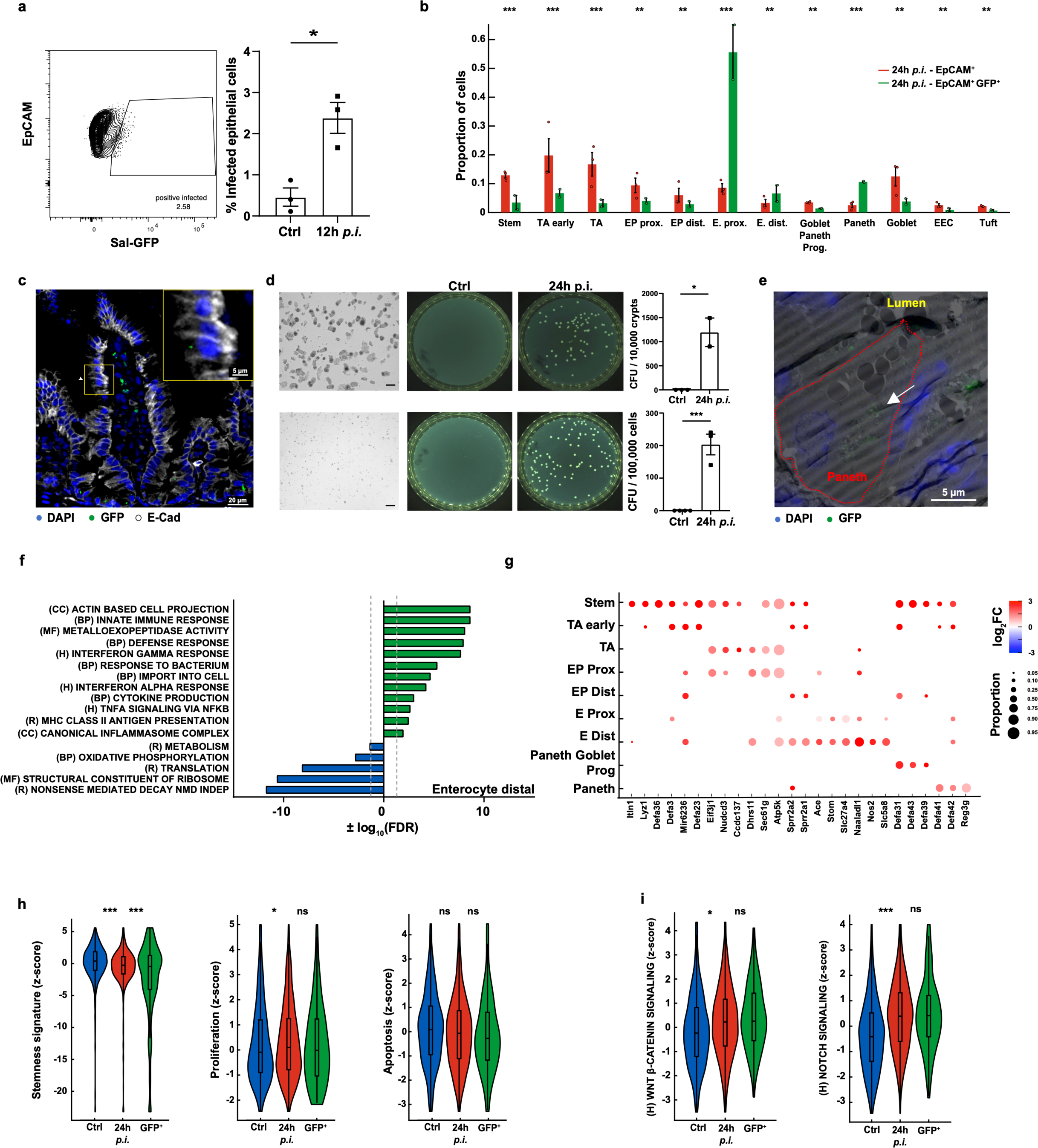
Salmonella specific-invasion signature of IECs. **a,** Representative flow cytometry plot (left) and quantification (right) of invaded cells (Sal-GFP^+^) gated on EpCAM^+^ cells 12h *p.i.. n* = 3 mice. Data are presented as mean ± SEM. \**p*-value < 0.05, two-tailed student’s *t-test*. **b,** Bar plots showing cell type proportions for each mouse (points) and condition; 24h *p.i.* EpCAM+ (red), and 24h *p.i.* EpCAM+ GFP+ (green) . Data are presented as mean ± SEM, statistical significance was determined by Poisson-glm (**Methods**), stars indicate FWER-Bonferroni adjusted \*\**p* < 2.5x10^-5^, \*\*\**p* < 2.5x10^-5^. **c,** Representative images of *Salmonella*-GFP staining from ileal villus at 24h *p.i..* IFA for *Salmonella*-GFP (green) and E-Cadherin (E-cad, white). Nuclei are shown using DAPI (blue). Scale bar, 20 µm. Top-right, Inset, 3x, showing *Salmonella* particle in distal enterocyte. Scale bar, 5 µm. **d,** Representative images of intestinal crypts (top left; scale bar, 100 µm) and crypt dissociated cells (bottom left; scale bar, 100 µm), accompanied by corresponding LB agar plates (middle panels, top and bottom) from control or 24h *p.i.*. CFU quantification (right panels) per 10,000 crypts (top) or 100,000 cells (bottom). Data are expressed as mean ± SEM from two independent experiments. \**p* < 0.05, \*\*\**p* < 0.001, two-tailed student’s *t-test*. **e,** CLEM image of Paneth cell infected with *S. enterica* (green) and nucleus (blue). Red dashed line, Paneth cell boundaries; white arrows indicate *Salmonella*-GFP particles. Scale bar, 5 µm. **f,** Selected gene sets showing hyper-geometric enrichment when testing the 400 significantly upregulated and downregulated genes in distal enterocytes 24h *p.i.* vs. control. H, Hallmark; R, Reactome; BP, GO biological process; CC, GO cellular compartment; MF, GO Molecular Function. **g,** Dot plot of significantly differentially expressed genes (DEG) across cell subsets comparing invaded (EpCAM+ GFP+) vs. infected (EpCAM+) cells from 24h *p.i.* infected mice. Size of the dot indicates the frequency of cells expressing the corresponding gene, while color indicates log2-fold change of mean expression in invaded (GFP+) cells. Only significantly DEGs are shown. **h,** Distribution of the expression of ‘Stemness’ (left), proliferation (middle) and apoptosis (right) gene signatures in control (ctrl), infected (24h *p.i.*) and invaded (GFP^+^) Lgr5+-ISC. Significance is determined by likelihood ratio test; ns, not significant; \**p* value < 0.05, \*\*\**p* value < 5x10^-5^. **i,** Distribution of the expression of Notch (left) and Wnt/b-catenin (right) gene signatures in control (ctrl), infected (24h *p.i.*) and invaded (GFP^+^) Lgr5+-ISC. Significance is determined by Mann–Whitney U test; ns, not significant; \**p* value < 0.05, \*\*\**p* value < 5x10-8.

**Extended Data Figure 3:**
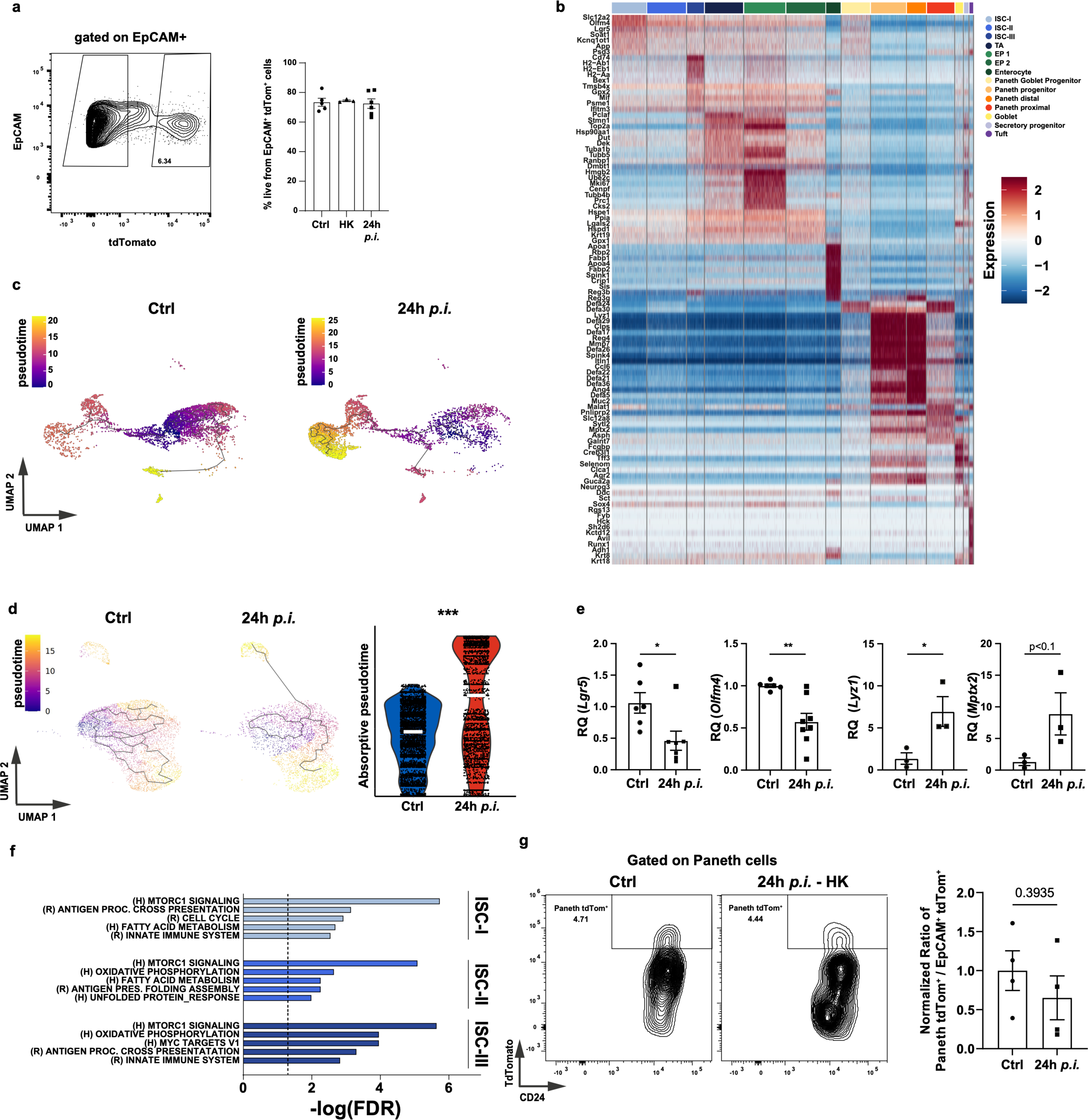
scPulse-seq of *Salmonella* infected IECs. **a,** Flow cytometry gating strategy of tdTomato+ IECs (EpCAM^+^ CD45^-^ Tom^+^). Indicated numbers are frequencies of tdTomato cells from the parental population (EpCAM^+^ CD45^-^). Frequency of live cells from EpCAM^+^ CD45^-^ Tom^+^ was analyzed (right, 3 mice per group); Data are presented as mean ± SEM; one-way ANOVA, ns, not significant. **b,** Cell subset signatures. The normalized expression level (row-wise z-score of scTransform normalized expression; color scale) of genes (rows) across cells (columns) is shown, sorted by cell subset. The cell subsets bar is on top, and color coding is shown on the top right. **c-d,** UMAP visualization of the secretory (**c**) or absorptive (**d**) tdTomato+ IECs superimposed with their pseudotime trajectories. Cells colored by pseudotime value, calculated with Monocole3 (scale per UMAP: top left; **Methods**). (**d**) Pseudotime distributions in absorptive path, binned by condition, comparing the differentiation state of control and 24h *p.i.* (right), **e,** Relative quantification (RQ) of Lgr5+-ISC markers, *Lgr5* and *Olfm4* (left) or Paneth cell markers *Lyz1* and *Mptx2* (right) in EpCAM^+^ Tom^+^ from control compared to 24h *p.i*. using quantitative PCR (qPCR). *n*≥3 mice per condition, \**p* value <0.05, \*\**p* value <0.01, two-tailed student’s *t-test.* **f,** Selected gene sets showing hyper-geometric enrichment when testing the 400 significantly upregulated genes in the three subsets of Lgr5+-ISC (ISC-I, II, III) 24h *p.i.* vs. control. H, Hallmark; R, Reactome. **g,** Representative flow cytometry plot (left) of the newly differentiated Paneth cells (tdTomato+, Paneth tdTom^+^) and quantification of the normalized ratio of Paneth tdTom^+^ compared to all tdTomato+ EpCAM+ (EpCAM^+^ tdTom^+^) cells in control and Heat-Killed (HK) *Salmonella* infected mice; *n* = 4 mice per condition. ns, non significant, two-tailed Student *t-test*.

**Extended Data Figure 4:**
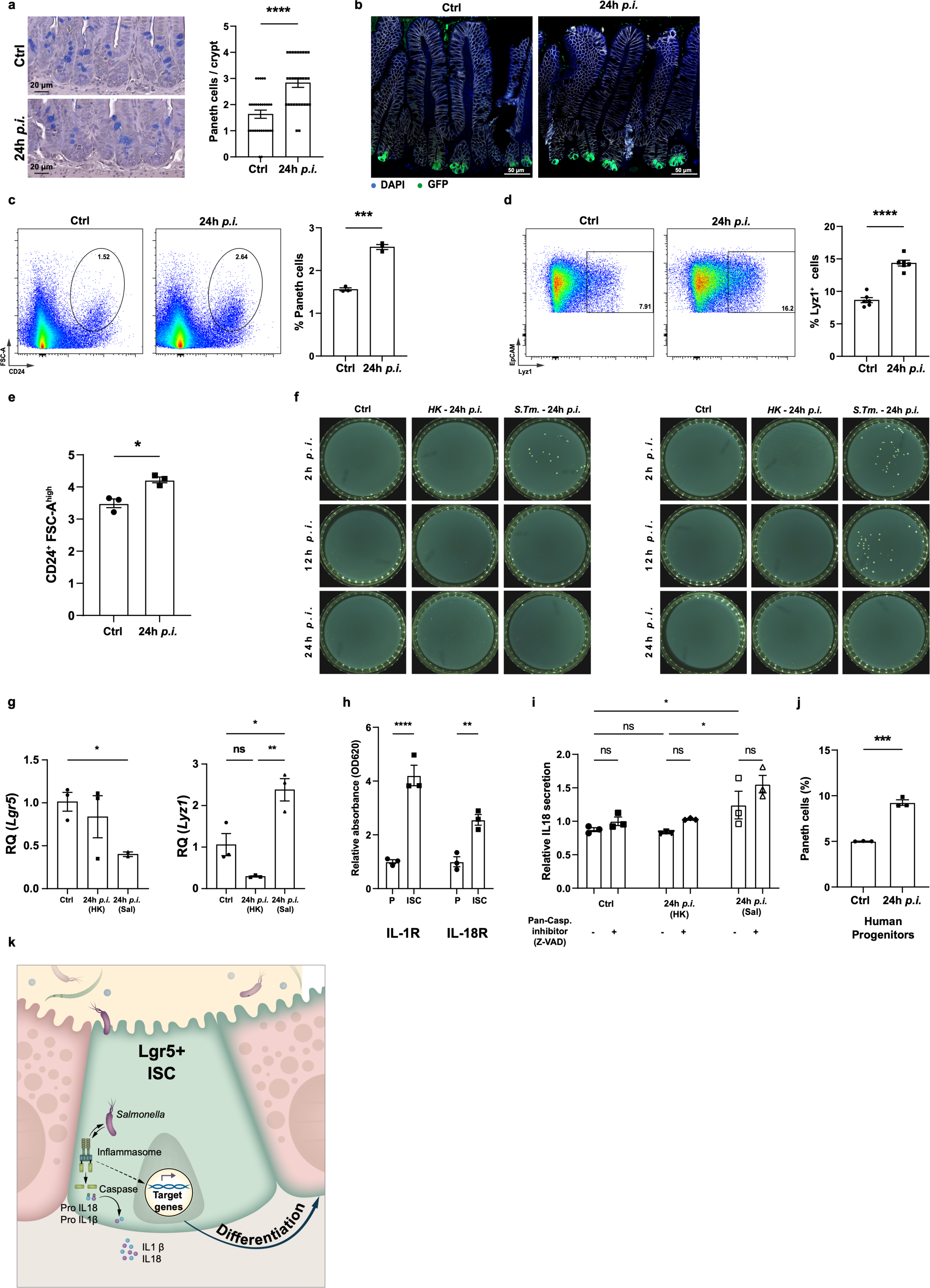
Stem cell responses to *Salmonella enterica* invasion. **a,** Periodic Acid Schiff (PAS) and Alcian Blue (AB) staining of ileum sections from control and 24h *p.i.* of TKO mice, scale 20μm. The number of Paneth cells per crypt was quantified (right panel); *n* ≤ 25 crypts per condition; data are presented as mean ± SEM, two-tailed student’s *t-test*, ****p-value < 0.0001. **b,** Distal SI section of infected TKO mice. IFA of Lyz1 (green) and E-Cadherin (E-cad, white). Nuclei are shown using DAPI (blue). Scale 50μm. **c-d,** Paneth cell number evaluated in organoids by CD24^+^ FSC^high^ (**c**) or Lyz1^+^ (**d**) gating (left). Frequencies of cells from control organoids (ctrl) or infected with *Salmonella*-RFP were quantified (right); *n* = 3 (c) or 6 (d) biological replicates per condition. Data are presented as mean ± SEM; \*\*\**p* value < 0.001, \*\*\*\**p* value < 0.0001, two-tailed student’s *t-test*. **e,** Frequencies of paneth cells (CD24^+^ FSC-A^high^) cells from control (ctrl) or *Salmonella*-RFP infected Lgr5-ISCs tested by flow cytometry. *n* = 3 biological repetitions. Data are presented as mean ± SEM; *p-value < 0.05, two-tailed student’s *t-test*. **f,** Representative *Salmonella* CFU. LB bacterial plates of *Salmonella* infected Lgr5-ISC or Paneth cells seeded after 2-, 12- or 24-h of *in vitro* infection. **g,** Relative quantification (RQ) of ISC marker, *Lgr5* (left) or Paneth cell marker *Lyz1* (right) of Lgr5+-ISC from control, or heat-killed (HK) or RFP *Salmonella* infected organoids, 24-hours post infection, using quantitative PCR (qPCR). Data are presented as mean ± SEM; ns, non significant, *p* value *<* 0.05, \*\**p* value < 0.01, one-way ANOVA. **h,** Relative absorbance of interleukin (IL)-1a,b or IL-18 in Lgr5+-ISC (ISC) or Paneth cells (P) media 12h *p.i..* Data are presented as mean ± SEM; \*\**p* value < 0.01, \*\*\**p* value < 0.001, two-tailed student’s *t-test*. **i,** Relative quantification of IL-18 secretion in media of control, heat-killed (HK) or *Salmonella*-RFP infected Lgr5+-ISC 24h *p.i.*. Data are presented as mean ± SEM; ns: non-significant, \**p* value < 0.05, one-way ANOVA. **j,** Frequencies of Paneth cells derived from control human intestinal progenitors cells (ctrl) or challenged with *Salmonella*-RFP (*n* = 3) tested by FACS. Data are presented as mean ± SEM \**p* value < 0.001, two-tailed Student’s *t-test*. **k,** Graphical abstract, novel stem cell defense mechanism mediated by inflammasome against intracellular pathogen, *Salmonella enterica*.

## Methods

### Mice

All mouse work was performed in accordance with the Institutional Animal Care and Use Committees (IACUC, no. 03240524-2) of the Weizmann Institute of Science. Mice were housed under specific-pathogen-free conditions at the animal facilities at the Weizmann Institute of Science, Rehovot, Israel. C57BL/6J wild type (WT) mice were purchased from Envigo. The following strains were obtained from the Jackson Laboratory: Lgr5-eGFP-IRES-CreERT2 (strain name: B6.129P2-Lgr5^tm1(cre/ERT2)Cle^/J; stock number 008875), Rosa26-LSL-tdTomato (strain name: B6.Cg-Gt(ROSA)26Sor^tm14(CAG-tdTomato)Hze^/J; stock number 007914), TKO (strain name: B6.129S-Rag2^tm1Fwa^ Cd47^tm1Fpl^ Il2rg^tm1Wjl^/J; stock number 025730). Lgr5-2A-eGFP reporter mice and Lgr5-2A-CreERT2 mice were gifts from Barker N. and were described before^85^. For lineage-tracing experiments, Lgr5-2A-CreERT2 mice were crossed to the Rosa26-LSL-tdTomato strain. ASC^ΔISC^ mice were generated by crossing ASC^loxp^ (ASC^flox/flox^) mice^86^ with Lgr5-2A-CreERT2 mice. Female age-matched mice from 7 to 10 weeks were used for all experiments in this study. Littermates of the same genotype and age were randomly assigned to experimental groups.

### Mice infection

Cultures of *S.Tm* were grown in Luria-Bertani (LB) medium at 37°C overnight to a stationary phase and were diluted 1:20 in fresh LB 4-hours prior infection. The cultures were washed in PBS and counted using OD600. Mice were deprived of food and water 4h prior infection. Gavage was performed in a volume of 100 μL containing either 5x10^8^ CFU of *Salmonella* strain SL1344 or PBS (as control). Mice were euthanized by CO_2_, 12-, 24-h post-infection. For all mice, ileal crypts were isolated from the small intestine as described below and used for subsequent experiments. Colony forming units (CFU) number was evaluated by plating serial 10-fold dilutions of homogenized crypts or IECs on selective Streptomycin (50μg/mL) or Ampicillin (100 µg/mL) agar plates.

### Lineage tracing and scPulse sequencing

To label Lgr5+-ISC and their following progeny for *in vivo* lineage tracing, we administered one intraperitoneal injection of Tamoxifen (2 mg per 20 g body weight) to induce Cre-mediated excision of a stop codon and subsequent expression of tdTomato from the Rosa26 locus. This injection occurred as soon as the mice received *per os S. Tm.* or PBS control.

### ASC deletion in intestinal epithelial cells

Cre activity was induced in 7-10 weeks old ASC floxed mice that expressed or not expressing Lgr5 driven CreER^T2^ by intraperitoneal injection (*ip*) of tamoxifen (Sigma-Aldrich), diluted in corn oil, 2 mg per injection, 5 times, every other day. Mice were sacrificed 10 days after the first injection. Organoids from these mice were used for subsequent experiments.

### Bacterial Strains

All *S.Tm* strains used in this study were derived from the wild-type strain SL1344, harboring a constitutive-GFP expressing plasmid (pFPV25.1; Addgene) or constitutive-RFP expressing plasmid (pC007; Addgene). *Salmonella*-GFP and *Salmonella*-RFP were provided by Prof. Roi Avraham from the Weizmann Institute of Science.

### Epithelial cell dissociation and crypt isolation

For all mice, crypts were isolated from the distal part of the SI. The SI was extracted and rinsed in cold Phosphate buffered saline (PBS). The tissue was opened longitudinally and sliced into small fragments roughly 2 mm long followed by incubation with 20mM EDTA-PBS on ice for 75 min. Then, the tissue was shaken vigorously, and the supernatant was collected as fraction 1 in a new conical tube. The tissue was incubated in fresh PBS, and a new fraction was collected every 5 min. Fractions were collected until the supernatant consisted almost entirely of crypts. The fractions 3-4 (enriched for crypts) were filtered through a 70 µm filter, centrifuged at 400g for 5 min, and dissociated with TrypLE Express (Gibco) for 1 min and 30 seconds at 37°C. The single-cell suspension was then passed through a 40μm filter and stained with fluorescence-activated cell sorting (FACS) antibodies and sorted with SH800 Sony sorter for subsequent analysis.

### Flow cytometry analysis

Single-cell suspensions were prepared as described above. Epithelial cells were stained with EpCAM (Biolegend), CD45 (Biolegend), CD24 (Biolegend), I-A/I-E (Biolegend) and Dapi-NucBlue (ThermoFisher) for 30’ on ice. Analysis was done on Cytoflex analyzer or Sony sorter. Further analysis of the data was done using FlowJo v10.1.

### Cell sorting

For the bulk population, FACS (SH800 Sony) was used to sort 1,000-5,000 cells into an Eppendorf tube containing 50ul TCL buffer (QIAGEN) solution with 1% 2-β-mercaptoethanol (Sigma-Aldrich). To enrich for Lgr5+-ISC population, cells were isolated from Lgr5-EGFP-CreER^T2^ or Lgr5-2A-CreER^T2^ mice, stained with the Dapi-NucBlue (ThermoFisher, cat no. R37605), CD45 (Biolegend, 30-F11), EpCAM (Biolegend, G8.8), CD24 (Biolegend, M1/69), Lyz1 (Dako, EC 3.2.1.17) and gated for EpCAM^+^ CD45^-^ (IEC), EpCAM^+^ CD45^-^ GFP^high^ (Lgr5+-ISC) or EpCAM^+^ CD45^-^ CD24^+^ FSC^high^ (Paneth cells). For bulk RNA-sequencing, the tubes were centrifuged and immediately frozen on dry ice and kept at 80°C until ready for the RNA isolation. For scRNA-sequencing, the cells were treated as described below.

### Hash-tagged single cell RNA-sequencing

Single cell RNA-sequencing (scRNA-seq) libraries were prepared using the chromium single cell RNA-seq platform (10x genomics). Isolated IEC were sorted into a cooled 15 mL tube with 0.04% BSA in PBS using a Sony SH800 cell sorter. Cells for each condition or mouse were stained using Biotin anti-mouse CD326 (EpCAM) antibody (cat #118204) followed by staining using TotalSeq™ PE Streptavidin (B0951 - B0955). 20,000 cells were sorted for each mouse or condition and pooled following manufacturer’s instructions. 30,000 single-cell suspension was loaded onto Next GEM Chip G targeting 15,000 cells and then ran on a Chromium Controller instrument to generate GEM emulsion. Single-cell 3’ RNA-seq libraries and cell surface protein libraries were generated according to the manufacturer’s protocol (Chromium Single Cell 3’ Reagent Kits User Guide (v3.1 Chemistry Dual Index). Final libraries were quantified using NEBNext Library Quant Kit for Illumina (NEB) and high sensitivity D5000/D1000 TapeStation (Agilent). Libraries were pooled according to targeted cell number, aiming for ∼20,000 reads per cell for gene expression libraries and ∼5,000 reads per cell for cell surface protein libraries. Pooled libraries were sequenced on a NovaSeq 6000 instrument using an S1 100 cycles reagent kit (Illumina).

### Bulk population RNA purification and cDNA preparation

Libraries were prepared using a modified SMART-Seq2 protocol^87^. RNA lysate cleanup was performed using RNAClean XP beads (Agencourt), followed by reverse transcription with Maxima Reverse Transcriptase (Life Technologies) and whole transcription amplification (WTA) with KAPA HotStart HIFI 2 3 ReadyMix (Kapa Biosystems) for 18 cycles. WTA products were purified with Ampure XP beads (Beckman Coulter), quantified with Qubit dsDNA HS Assay Kit (ThermoFisher). The products were diluted into a concentration of 1 ng/μL in ultra-pure water.

### Quantitative PCR

For quantitative PCR (qPCR), cDNA from bulk population libraries were diluted to a final concentration of 0.05 ng/μL. Fast SYBR green master mix (Thermo-Fisher scientific) was used to perform qRT–PCR per the manufacturer’s instructions. The primer sequences are used are: Lgr5 Forward: 5’ – GACAATGCTCTCACAGAC – 3’; Lgr5 Reversed: 5’ – GGAGTGGATTCTATTATTATGG – 3’; Smoc2 Forward: 5’ – GGAGCAGGGAAAGCAGATGAT – 3’; Smoc2 Reverse: 5’ – GGTCTTGTTCTGCCGACTCT – 3’; OLFM4 Forward: 5’ – CACAGCTCACATCCTTTCTCAG – 3’ ; OLFM4 Reverse: 5’ – ACTCGGACCGTCAGGTTCAG – 3’; Lyz1 Forward: 5’ – CAAAGAGGGTGGTGAGAGATC – 3’; Lyz1 Reversed: 5’ – TGAGAAAGAGACAGAATGGGC – 3’; Mptx2 Forward: 5’ – CTCTCTGTTCTTTCAGGAAGTGTAGC – 3’; Mptx2 Reversed: 5’ – ACACATAGGCAGTGGATGATTCTT – 3’; UBC Forward: 5’ – AACATCCAGAAAGAGTCCACC – 3’; UBC Reversed: 5’ – CATTCTCTATGGTGTCACTGGG – 3’. Data analysis was performed with QuantStudio 12K flex software (Thermo-Fisher scientific) based on the ΔΔCT method. All target genes were standardized to endogenous reference gene Ubiquitin C (*UBC*).

### Histochemistry

Tissues were fixed for 16-24 hours in formalin, embedded in paraffin, and cut into 5μm thick sections. Sections were deparaffinized with standard techniques. For H&E staining, slides were stained with Hematoxylin for 1 minute, washed, stained with Eosin for 45 seconds. For PAS and Alcian blue staining, tissue was stained with Alcian blue and periodic acid-Schiff reagents kit (ScyTek Laboratories, cat no. APS-1) on deparaffinized slides with standard techniques, using 3% acetic acid for 3 minutes, then Alcian blue solution (Alcian blue, pH 2.5) for 30 min, rinsed in tap water for 5 min, and oxidized in periodic acid (0.5%) for 10 min, followed by rinsed in running tap water for 5 min, and stained in Schiff reagent as a counterstain (cancer diagnostics) for 10 min. Nuclei were stained with PureView™ Mayers Hematoxylin for 50 seconds. All slides were washed, dehydrated and mounted with Sub-x mounting medium (Leica).

### Immunofluorescence (IF)

Sections were deparaffinized with standard techniques and incubated with primary antibodies overnight at 4°C, followed by secondary antibodies incubation at room temperature for 45 min. All slides were mounted with Slowfade Mountant+DAPI (Life Technologies, S36964) and sealed. Antibodies used for immunofluorescence were mouse anti-E-cadherin (1:100, BD Biosciences Clone 36), rabbit anti-Olfm4 (1:200, Cell Signaling Technology D6Y5A), rat anti-Ki67 (1:100, Invitrogen SolA15), rat anti-lysozyme (1:200, Dako A0099), rabbit anti-cleaved-Caspase 3 (1:100, Cell Signaling Technology 9661S), rabbit anti-GFP (1:100, Cell Signaling Technology 2956S) and Alexa Fluor 488-, 568- and 647-conjugated secondary antibodies (1:400, Abcam).

### TUNEL assay

Small intestine tissue was analyzed for TUNEL staining by the TUNEL Assay Kit (Abcam cat no. ab66110) according to the manufacturer’s recommendations. In brief, sections were deparaffinized followed by antigen retrieval, labeled with BrdU for 1 hour at 37°C, and incubated with anti– BrdU-Red antibody for 30 minutes at room temperature. After TUNEL labeling, the sections were stained with E-cadherin antibody (Biolegend cat no. 118218) then washed and mounted with Slowfade Mountant+DAPI (Life Technologies, S36964). Measurement and analysis of the staining area was performed using Zen software (Zeiss).

### Single molecule fluorescence in situ hybridization (smFISH)

A RNAScope Multiplex Fluorescent Kit (Advanced Cell Diagnostics) was used as per the manufacturer’s recommendations with the following alterations. The target retrieval boiling time was adjusted to 14 min and incubation with Protease IV at 40 °C was adjusted to 8 min. Slides were mounted with Slowfade Mountant+DAPI (Life Technologies, S36964) and sealed. Probes used for single-molecule RNAscope (Advanced Cell Diagnostics): *S.typhimurium*-16SrRNA-O1 (C1) and *Lgr5* (C2).

### Combined immunofluorescence and smFISH

Following the Amp 4 step of the smFISH protocol as described above, tissue sections were washed and incubated overnight at 4°C with primary antibodies, washed three times in 1× TBST, and then incubated with secondary antibodies for 45 minutes at room temperature. Slides were then mounted with Slowfade Mountant + DAPI (Life Technologies, S36964) and sealed.

### Image analysis

Images of tissue sections were taken with a confocal microscope LSM900 confocal microscope (Zeiss). Scale bars were added to each image using the confocal Zen analysis software (Zeiss). Images were overlaid and visualized using Zen analysis software (Zeiss). Organoids and single-cell suspension were imaged using a bright field optical microscopy (EVOS™ M5000 Imaging System) and analyzed with FIJI software.

### Image quantification

Quantification of IFA images of all tissues were assessed by staining for E-Cadherin (BD Biosciences) to mark cell borders, and DAPI staining for nucleus visualization. Cells were manually counted based on immunofluorescence staining specific to each cell type. Stem cells were quantified using the ISC marker Olfm4 (Cell Signaling Technologies) within the “stem cell zone” at the bottom of the crypt, delineated by a dashed line, with quantification restricted to this area. Paneth cells were similarly quantified using Lyz1 (Dako) as the marker. Proliferative cells (Ki67^+^) were counted along the villus-crypt axis. For each quantification, at least fifteen intact and longitudinally oriented crypts per tissue section were analyzed.

### Transmission electron microscopy

Samples were fixed in 4% paraformaldehyde with 0.1% glutaraldehyde in 0.1M cacodylate buffer (pH = 7.4) for 1 hour at room temperature and kept overnight at 4° C. The samples were soaked overnight in 2.3M sucrose and rapidly frozen in liquid nitrogen. Frozen ultrathin (70–90 nm) sections were cut with a diamond knife at −120°C on a Leica EM UC6 ultramicrotome. The sections were collected on 200-mesh Formvar coated nickel grids. Sections were blocked with a solution containing 1% BSA, 0.1% glycine, 0.1% gelatin, and 1% Tween 20. Contrast staining and embedding were performed as previously described (Tokuyasu, 1986). The embedded sections were viewed and photographed with a FEI Tecnai SPIRIT (FEI, Eidhoven, Netherlands) transmission electron microscope operated at 120 kV, and equipped with a OneView Camera.

### Correlative light and electron microscopy

Wide-field fluorescence images of the thin sections on fresh sliced TEM grids (before contrast staining) were taken using VUTARA SR352 system (Bruker) with 1.3 NA 60x silicon oil immersion objective (Olympus). Z slices of 150 nm were collected using 405 nm and 488 nm excitation lasers in the presence of a buffer containing 7 μM glucose oxidase (Sigma), 56 nM catalase (Sigma), 2 mM cysteamine (Sigma), 50 mM Tris, 10 mM NaCl, 10% glucose, pH 8. Contrast staining and embedding were performed as previously described (Tokuyasu, 1986). The embedded sections were viewed and photographed with a FEI Tecnai SPIRIT (FEI, Eidhoven, Netherlands) transmission electron microscope operated at 120 kV, and equipped with a OneView Camera. Overlay TEM and Fluorescence images was done using nuclei marked with DAPI as fiducial markers using Photoshop.

### *In vitro* infection

Following crypt isolation from different mouse models (WT, Lgr5-CreER^T2^-EGFP, ASC^ΔISC^), crypts were resuspended in 500 μL of L-WRN conditionated media (C.M.)^61^ supplemented with 10 µM Y-27632 Dihydrochloride (Biogems, cat. no 1293823). The number of crypts was estimated using a bright field optical microscopy (EVOS™ M5000 Imaging System). The crypts were incubated with either L-WRN C.M. or L-WRN C.M. containing *S. Typhimurium* RFP (MOI = 10, about 5.10^5^ CFU / 500 crypts) for 2-hours at 37 °C. After 3 washes with DMEM/F-12 containing 100 µg/mL gentamicin, about 500 crypts were embedded in 20 μL Matrigel™ (Corning) with 1µM Jagged-1 peptide (Ana-Spec). After 20 min at 37 °C in a humidified atmosphere at 5% CO_2_, L-WRN C.M. supplemented with 10 µM Y27632 were added and incubated at 37 °C with 5% CO_2_ for the time of the experiment. For experiments requiring pan-caspase inhibition, Z-VAD-FMK (10 μg/ml; 20 μM) was added to the media.

#### Heat-killed

The same concentration of heat-killed bacteria was prepared by centrifuging bacteria and wash in PBS, followed by incubation at 95°C for 30 minutes in the desired concentration in PBS. A sample of HK *Salmonella* was seeded onto LB agar plates with streptomycin to confirm bacterial killing. After 2-, 12- or 24-hours, supernatants were used for HEK-Blue^TM^ IL-1R assay. Pellets containing cells or organoids were used for RNA isolation, flow cytometry analyses or CFU evaluation.

#### CFU evaluation

Cells or organoids were collected and incubated in 100 μL DDW with 0.2% Triton X-100 (Sigma-Aldrich, Israel) to release intracellular bacteria. After 2 minutes of incubation, tubes were spun down, resuspended in 100 μL of PBS and each tube was plated on selective Streptomycin (50μg/mL) or Ampicillin (100 µg/mL) agar plates. The plates were incubated overnight and the CFU per condition were counted manually.

#### HEK-Blue^TM^ IL-1R assay

IL-1R signaling activity was assessed using HEK-Blue™ IL-1R cells (Invivogen, cat. code hkb-il1r) according to the manufacturer’s protocol. HEK-Blue^TM^ IL-1R cells were cultured in DMEM supplemented with 10% heat-inactivated FBS, and 100 µg/mL Normocin. Cell lines were cultured at 37 °C and 5% CO_2_ and routinely tested for mycoplasma contamination using a MycoBlue Mycoplasma Detector kit (Vazyme Biotech, cat: D101-01). For the assay, the supernatant from organoids, Lgr5+-ISC or Paneth cells was centrifuged at 1000 × g for 5 minutes to eliminate cell debris. A 20μl aliquot of the supernatant was added to 5 × 10^4^ HEK-Blue™ IL-1R cells seeded in a 96-well plate. Complete medium, with or without recombinant IL-1β (0.1 – 1000 pg/ml) (Peprotech, cat. no. 211-11B), served as positive and negative controls, respectively. The following day, 20μl of the HEK cell supernatant was transferred to a flat-bottom 96-well plate containing 180μl of QuantiBlue (Invivogen, cat. code rep-qbs) detection reagent and incubated at 37°C for 2 hours. IL-1R activity was measured using a spectrophotometer (Tecan) at 630 nm. Standard curves were performed using dilutions of murine IL-1β (0.1 – 1000 pg/mL).

#### Human organoid assay

Human ileal organoids used in this study were obtained from the Israeli Hadassah Organoid Center (HOC) and generated from biopsies collected from the distal ileum. The collection and use of these organoids were approved under the Committee on Research Involving Human Subjects application (0921-20-HMO, The Israeli Organoid Bank- A Means For- Personalised Medicine) and the Weizmann Institute of Science IRB.

In brief, organoids were cultured in IntestiCult Organoid Growth Medium (Human, Stemcell, Cat no. 06010) in 3D Matrigel in 24-well plates. In experiments requiring differentiation, IntestiCult Differentiation Medium was used (Human, Stemcell, Cat no. 100-0214). Organoids were dissociated and infected with *S.Tm.* for two hours at 37°C. After 3 washes with DMEM/F-12 containing 100 µg/mL gentamicin, epithelial aggregates were embedded in 20 μL Matrigel™ (Corning) with 1µM Jagged-1 peptide (Ana-Spec). After 20 min at 37°C, the differentiation medium was added and incubated at 37 °C with 5% CO_2_ for the experiment.

For progenitor infection, after 2 day of growth in IntestiCult Orgnaoids Growth Media, organoids were dissociated using TrypLE Express (Gibco) for 1 min and 30 seconds at 37°C. The single-cell suspension was then passed through a 40μm filter and infected with *S.Tm.* for two hours at 37°C in suspension. After 3 washes with DMEM/F-12 containing 100 µg/mL gentamicin, cells were left in suspension in differentiation media at 37°C for 24h.

### Statistical analysis

Statistical analysis was performed with the Prism software (GraphPad software), R v4.4.1, R v4.3.1, and Matlab R2020a. The specific tests applied are detailed in the corresponding figure legends and computational methods.

## Computational methods

### scRNA-seq pre-processing and demultiplexing

For droplet-based scRNA-seq, CellRanger v7 was used to align reads to the GRCm39 (m32) mouse genome reference. In a subset of batches multiple (3 or 4) experimental conditions were overloaded onto a 10x flow channel^88^ and demuxing was performed by Pegasus demuxEM (v1.3.1) using default settings (https://github.com/lilab-bcb/demuxEM/^89^).

### scRNA-seq processing, quality control filtering

scRNAseq reads were re-aligned with STARsolo to the GRCm39 (m32) using multimapping mode, quantifying both full-length and exonic-only transcripts (v2.7.10)^90^. The output was processed using the dropletUtils R package (version 1.7.1), to exclude any chimeric reads that had less than 80% assignment to a cell barcode, and identify and exclude empty cell droplets^91,92^, by testing against a background generated from barcodes with 1,000 to 10 UMIs, with cutoffs determined dynamically based on channel-specific characteristics. Next, we used Cellbender (^93^, v3.0), to exclude ambient transcripts and quantify the ambient proportion in each cell barcode. Cell barcodes were excluded if they satisfied any one of the following criteria: (1) Fewer than 300 genes; (2) Fewer than 500 UMIs (after excluding MT and intron UMIs); (5) Non-empty droplet with false discovery rate (FDR) less than 0.1; or (6) Over 30% of reads estimated as coming from ambient reads according to cellbender.

Using outlier exclusion separately for each each 10x channel, cells that deviated by > 2 interquartile ranges (IQR) from the median were then flagged based on the following criteria: (1) log10(total transcript UMI), (2) Fraction of barcode swaps, (3) Gene saturation estimate, (4) UMI saturation estimate, (5) Fraction of UMI supported by > 1 read^94^. Cells were further flagged if they substantially deviated from the fit based on the following relationships: (1) Total reads versus total UMI, (2) Total UMI versus log likelihood of being empty^92^; (3) Total UMI versus total number of genes. A cell was excluded if it was flagged by at least two of these criteria.

### Selection of variable genes, dimensionality reduction, clustering and cell-type assignment

After quality filtering and exclusion, scRNA-seq expression profiles were clustered using a non-negative matrix factorization (NMF)^95^ and a graph clustering-based approach. Transcriptionally over-dispersed genes were identified within each experimental batch (i.e., 10x channel) by the difference of the coefficient of variation (CV) from the median CV for other genes with a similar mean expression^96^. A robust set of 1,000 to 6,000 genes was retained based on an elbow-based criterion, applied to the median of over-dispersed difference statistics based on 200 samples of 75% of cells.

In subsequent analysis of single cell data was sequencing depth normalized to log2(TP10K+1) values. Next, 80% of genes and samples were sub-sampled between 100 times, and NMF was used to reduce the dimensionality of the full dataset to between 15 and 40 dimensions as the product of two non-negative matrices^97^. The loading matrices (i.e., activations) of these NMF components were used to calculate the k-nearest neighbors (k-NN) graph (k = 21) based on a cosine similarity distance. This graph was clustered using stability optimizing graph clustering (http://michaelschaub.github.io/PartitionStability/^98,99^), to identify cell type clusters. Cell types were manually annotated based on known cell-type markers^4^.

To minimize differences across samples due to technical reasons, gene expression measurements of individual genes were quantile normalized such that the expression CDFs for each gene matched.

Gradient boosting (R 3.6.1, xgboost v0.90.0.2 (https://doi.org/10.1145/2939672.2939785)) was first applied to build a cell to cluster classifier for each cluster type based on uninfected control. For each of the 12 cell types, a separate classifier was trained to predict each cell type separately (one-versusall), in a 5-fold cross-validation scheme. Next, using the probability scores of the held-out test-set we identified an optimal cutoff for each class based on an ROC analysis comparing the true positive rate (TPR = true positives divided by all positive predictions) to the false positive rate (FPR = true negative divided by all negatives) and selecting the point at which the ROC curve intersects with the diagonal. In cases where a cell was assigned to more than one subtype, we used the assignment with the higher predictive score. Cells that could not be assigned confidently by any classifier were excluded from further analysis.

### Visualization of single cell profiles

We generated uniform manifold projection maps (UMAPs), per set of experiments from consensus NMF loading matrices, with n_nieghbors value of 30, and min_dist value of 0.3, and euclidean distance.

### Identification of differentially expressed genes

Differentially expressed genes (DEGs) were identified using a negative binomial mixed model (NEBULA v1.5.2^100^, https://github.com/lhe17/nebula), with experimental batch as a random effect, an offset of total exonic-UMI counts per cell, and using an NBGMM model. DEG testing was performed in several ways and included in the supplementary tables:

(1) Global tissue test, *Sal24h p.i.* vs *uninfected controls* (**Supplementary Table 2**).
(2) Global ‘matched-tissue’, *Sal24h p.i.* vs *uninfected controls* (**Supplementary Table 2**).

Here, tissue was matched by subsampling cells such that the conditions are comprised of similar cell-type proportions. We report the trimean statistics over 30 random sub-samples.

(3) *Sal24h p.i.* vs *uninfected controls* (**Supplementary Table 3**). Performed separately per cell type.
(4) *Sal24h GFP+ p.i.* vs *Sal24h p.i* (**Supplementary Table 5**). Performed separately per cell type.
(5) *Tomato+ Sal+ p.i.* vs *Tomato+ Sal-* (**Supplementary Table 7**). Performed separately per cell type.

Genes were considered DEG if they passed the following criteria:

1. Nebula test successfully converged.
2. Nebula p-values after Benjamini-Hocheberg correction had FDR<0.1
3. Nebula log2 fold-change was > log2(1.5)
4. Gene showed non-zero expression in > 5%
5. Log-2 fold-change of the mean of expressing cells between conditions > log2(1.5)

### Gene-set enrichment of differentially expressed genes

To explore associations between differentially expressed genes (DEGs) and known gene sets, we implemented a classical gene-set enrichment analysis using a hypergeometric test. We selected the top 400 DEGs for comparison against a comprehensive set of 13,239 gene sets derived from the MisgDB compendium, which includes databases such as Biocarta, Reactome, WikiPathways, GTRD, Gene Ontology (GO), Molecular Signatures Database (MPT), and Hallmarks^101^. This analysis was further extended by incorporating bioPlanet genesets^102^ and additional gene sets identified in the current study based on the top DEGs. A geneset was considered significantly enriched if it contained at least four overlapping genes with our DEGs, and the hypergeometric test revealed enrichment with a false discovery rate (FDR) less than 0.1, adjusted using the Benjamini-Hochberg method.

### Calculating gene signature scores

Single-cell gene-set signature enrichment was performed as previously described^103^. Genes were categorized into 20 bins based on their mean expression across all cells, with bin boundaries determined by the overall gene expression distribution. Gene expression values were centered, scaled, and transformed to the [0,1] range using a logistic function. For a gene set signature comprising k genes, a raw signature score was calculated for each cell as the mean of the normalized expression values. This raw score was then compared to a null distribution generated from 1,000 randomly selected gene sets, each containing k genes. The random gene sets were constructed to match the global mean expression bins of the original signature genes.

The final per-cell activity score was derived by centering the raw score against the mean of the null distribution scores. For visualization, these activity scores were further standardized by median-centering and scaling by the mean absolute deviation (MAD). Significance between conditions was tested using a generalized linear mixed model with batch as a random effect and condition as the fixed effect. We report the p-values of a likelihood-ratio test comparing a baseline model with the model including the condition (fixed effect).

### Comparing cell-type proportions in single-cell experiments

The abundance of each cell type across analyzed mice was modeled as a Poisson-distributed random variable. We employed a generalized linear model (GLM) with the following specifications, the distribution family was ‘poisson’, response variable was the count of a tested cell type, offset variable as the total number of post-qc filtering profiled cells per mouse/batch, and predictor variable as the tested condition (*e.g. Sal 24h pi vs uninfected control*). The statistical significance of the treatment effect was evaluated using a Wald test on the corresponding regression coefficient.

### Single-cell signature in human data

We further validated our identified signature of top differentially expressed genes (DEGs) from the TomSal+ versus TomSal-experiment in a contemporary study on human inflammatory bowel disease (IBD) patients^50^. We selected genes consistently recognized as DEGs across three or more cell types, forming distinct positive and negative gene signatures, comprising 81 and 85 genes, respectively. These gene signatures were translated to human gene symbols through ortholog mapping^104^ and evaluated in human single-cell RNA sequencing (scRNA-seq) datasets, as detailed in our gene signature scoring methodology (see “Calculating Gene Signature Scores”). Signature scores were normalized using z-scores based on median and median absolute deviation (MAD), after which the negative signature scores were subtracted from the positive scores. Statistical significance was assessed using a likelihood-ratio test to compare a baseline model without the condition (fixed effect) against a model that includes it, with both models accounting for batch and chemistry as random effects.

